# Polycysteine-encoding leaderless short ORFs function as cysteine-responsive attenuators of operonic gene expression in mycobacteria

**DOI:** 10.1101/834739

**Authors:** Jill G. Canestrari, Erica Lasek-Nesselquist, Ashutosh Upadhyay, Martina Rofaeil, Matthew M. Champion, Joseph T. Wade, Keith M. Derbyshire, Todd A. Gray

## Abstract

Genome-wide transcriptomic analyses have revealed abundant expressed short open reading frames (ORFs) in bacteria. Whether these short ORFs, or the small proteins they encode, are functional remains an open question. One quarter of mycobacterial mRNAs are leaderless, beginning with a 5’-AUG or GUG initiation codon. Leaderless mRNAs often encode unannotated short ORFs as the first gene of a polycistronic transcript. Here we show that polycysteine-encoding leaderless short ORFs function as cysteine-responsive attenuators of operonic gene expression. Detailed mutational analysis shows that one polycysteine short ORF controls expression of the downstream genes. Our data indicate that ribosomes stalled in the polycysteine tract block mRNA structures that otherwise sequester the ribosome-binding site of the 3’gene. We assessed endogenous proteomic responses to cysteine limitation in *Mycobacterium smegmatis* using mass spectrometry. Six cysteine metabolic loci having unannotated polycysteine-encoding leaderless short ORF architectures responded to cysteine limitation, revealing widespread cysteine-responsive attenuation in mycobacteria. Individual leaderless short ORFs confer independent operon-level control, while their shared dependence on cysteine ensures a collective response mediated by ribosome pausing. We propose the term ribulon to classify ribosome-directed regulons. Regulon-level coordination by ribosomes on sensory short ORFs illustrates one utility of the many unannotated short ORFs expressed in bacterial genomes.

## INTRODUCTION

Short open reading frames (sORFs) are extremely difficult to computationally identify in genome sequences; their shortened gene length approaches statistical random ORF background frequencies and their amino acid sequences have limited bioinformatic value (Frith et al., 2006, Crappe et al., 2013, Hemm et al., 2008, Hobbs et al., 2011). Conventional mass spectrometry proteomic studies also systematically underrepresent the small proteins (sproteins) encoded by sORFs, as they can be lost during sample preparation or provide too few detectable peptides (Hemm et al., 2010). These limitations have contributed to both a lag in (i) recognition of sORFs, and (ii) assessment of their functional potential. This knowledge gap has been underscored by recent descriptions of many potential novel sORFs in *Escherichia coli* and in bacteria found in the human microbiome (Weaver et al., 2019, Meydan et al., 2019, Sberro et al., 2019). Previous work applying complementary transcriptomic approaches to mycobacteria, in both slow-growing *Mycobacterium tuberculosis*, and fast-growing *Mycobacterium smegmatis* and *Mycobacterium abscessus*, suggested that their genomes contained hundreds of sORFs actively producing sproteins (Miranda-CasoLuengo et al., 2016, Shell et al., 2015). Ribosome profiling (Ribo-seq), together with RNA-seq and transcription start site (TSS) mapping, provided genome-wide empirical evidence for the location and ribosome occupancy of mycobacterial mRNAs that express sproteins (Cortes et al., 2013, Shell et al., 2015, Miranda-CasoLuengo et al., 2016).

Leaderless translation is a phenomenon in which mRNAs without a 5’ UTR are efficiently translated, such that the first nucleotide of the start codon is the 5’ end of the RNA. We and others have shown that leaderless translation is abundant in mycobacteria (Cortes et al., 2013, Shell et al., 2015). Many mycobacterial ORFs that are predicted to be initiated by leaderless mRNAs (LL-mRNAs) are short (defined here as less than 150 nt) and unannotated. The ORFs initiated at the combined transcription/translation initiation start sites of LL-mRNAs are readily identifiable from transcriptomic data sets and thus provide a high-confidence list of expressed novel mycobacterial sORFs. Insights into the mechanistic and functional attributes of LL-mRNAs have lagged, as they are rare or poorly expressed in *E. coli* but are abundant in other bacteria, archaea, and mitochondria (Beck and Moll, 2018). LL-sORFs represent the first (5’-most) ORF of a transcript, which positions them for a potential role in *cis*-regulation of the downstream operonic genes. There is ample precedent in eukaryotes for regulation of downstream genes by such “upstream ORFs” (uORFs) (Couso and Patraquim, 2017, Hinnebusch et al., 2016). In prokaryotes, mechanisms have also been previously described in which uORFs have been shown to regulate expression of downstream genes through a process known as attenuation (Bechhofer, 1990, Henkin and Yanofsky, 2002, Oppenheim and Yanofsky, 1980).

Attenuation is a *cis*-regulatory mechanism often mediated by bacterial short uORFs enriched in codons for the amino acid product of that biosynthetic operon. Attenuation occurs when abundant charged tRNA levels allow translating ribosomes to quickly pass through the modulating uORF, promoting the formation of an intrinsic terminator that aborts transcription of the remainder of the operon. Low levels of charged tRNA cause ribosome stalling in the uORF at codons for the end-product amino acid, facilitating the formation of a competing anti-terminator structure, thereby releasing attenuation to allow transcription to extend into the biosynthetic operon (Turnbough, 2019)(Fig 1). To our knowledge, uORF-mediated attenuation mechanisms for cysteine have not been described. A subset of predicted mycobacterial LL-sORFs conspicuously encode consecutive cysteine residues, and these were found upstream of genes annotated to be involved in cysteine biosynthesis (Shell et al., 2015). We hypothesized that these LL-sORFs function as cysteine-sensitive attenuators.

**Figure 1.**
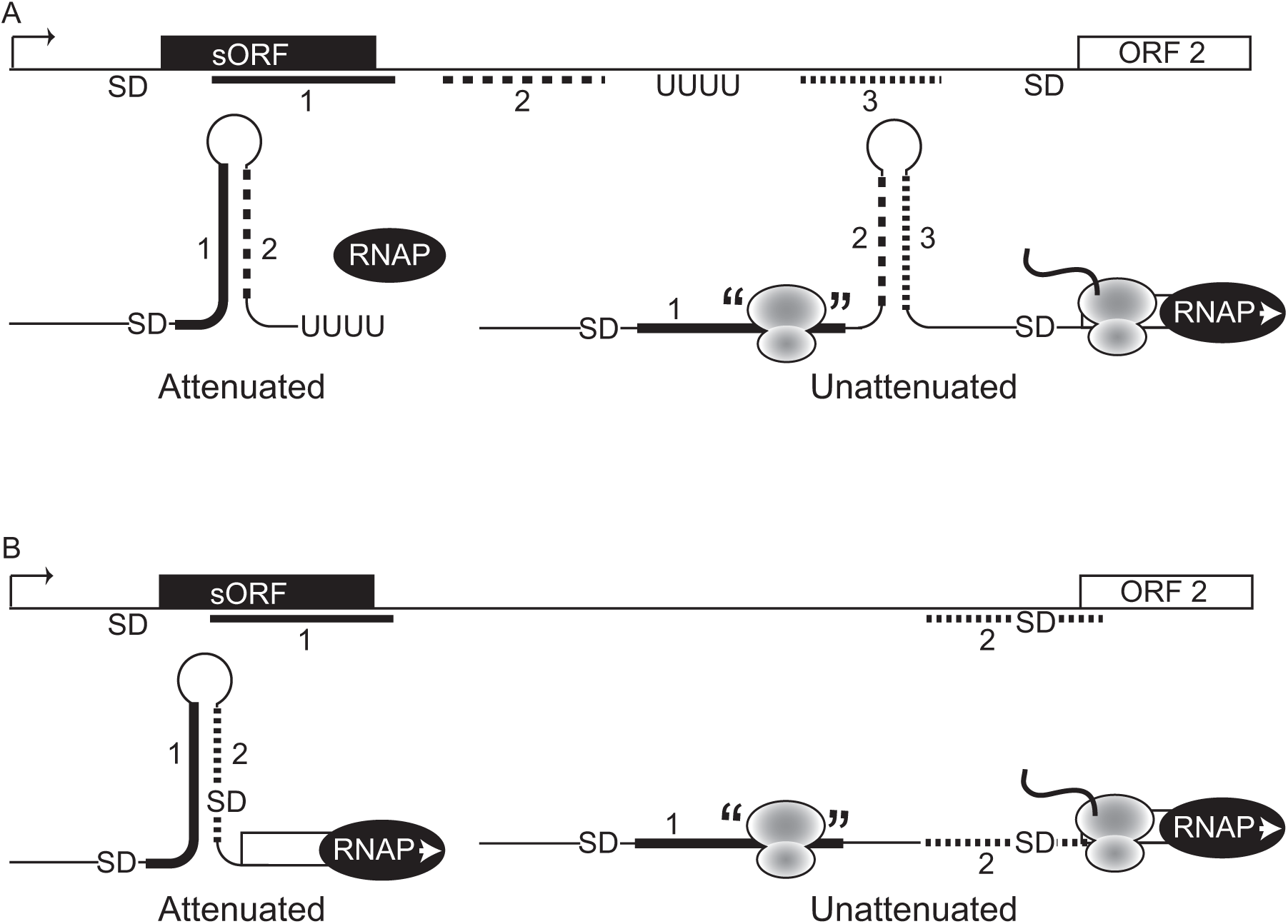
Schematic comparison of structural features of operon attenuation. A. In transcriptional attenuation, the balance of dueling mRNA hairpin structures is controlled by ribosome occupancy of the sensory sORF. Rapid ribosome transit--when tRNA^trp^ is plentiful, for example--allows the vacated sORF mRNA to hybridize with complementary sequence thereby forming the stem of an intrinsic transcriptional terminator. This transcriptional termination keeps the downstream tryptophan biosynthetic genes from unneeded expression. Under conditions of reduced tRNA^trp^ levels, ribosome pausing in the sORF blocks the attenuating transcriptional terminator from forming, an anti-terminator structure forms instead, which promotes RNA polymerase extension into the operon. Transcriptional attenuation through controlled intrinsic termination occurs in *E.coli* tryptophan and *S. typhimurium* histidine biosynthetic loci, and the whiB7 locus in *Mycobacterium abscessus* (Burian and Thompson, 2018, Johnston et al., 1980, Yanofsky, 1981). B. In translational attenuation, sensory sORF mRNA base pairing obscures the SD of the downstream gene, preventing translation initiation. Ribosome arrest in the sORF prevents duplex formation, and consequently frees the SD to recruit and position translation initiation complexes for expression of the annotated gene. Translation attenuation has been described for some ribosome antibiotics (Arenz et al., 2014, Lovett and Rogers, 1996, Ito and Chiba, 2013).

## RESULTS

We previously showed that an AUG or GUG (collectively, RUG) at the start of an mRNA will direct translation of an ORF determined by that initiation codon (Shell et al., 2015). Therefore, transcription start sites (TSS) beginning with RUG in mycobacteria both readily identify leaderless mRNAs, and also establish the resulting translated reading frames. Using published transcription start site mapping and ribosome profiling fata, we identified 304 putative LL-sORFs in *Mycobacterium smegmatis* (Table S1) (Martini et al., 2019, Shell et al., 2015). Cysteines that are encoded by consecutive codons (polycysteine) are particularly enriched in LL-sORFs when compared to annotated ORFs (Fig 2, Table S2). This enrichment of polycysteine in LL-sORFs is reminiscent of consecutive tryptophans or histidines in leader sORFs in classic attenuated operon models (Yanofsky, 1981).

**Figure 2.**
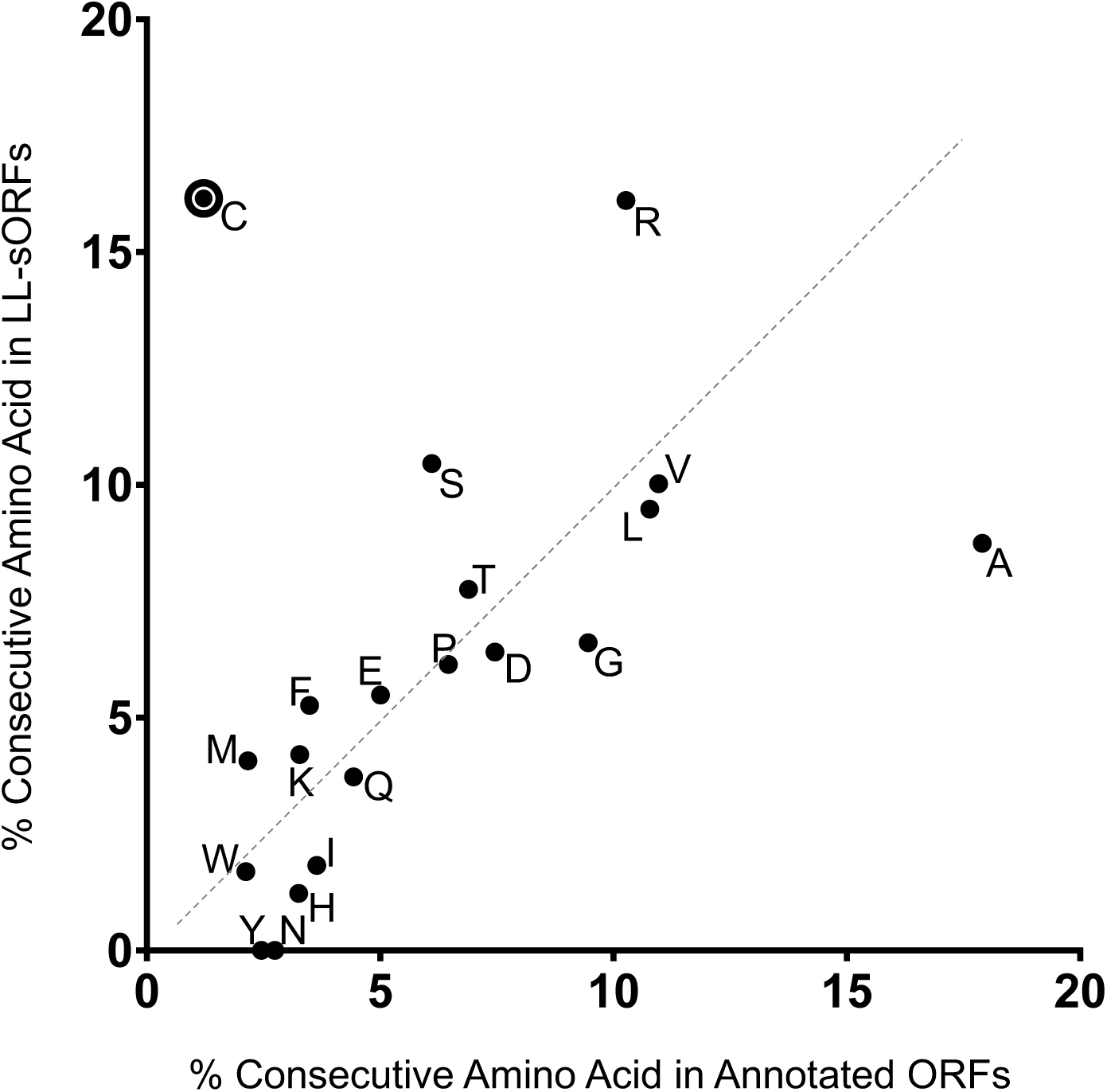
Polycysteines are enriched in LL-sORFs. The frequency of consecutive amino acids is shown in annotated genes (X-axis) and in LL-sORFs (Y-axis). Frequencies for each clustered amino acid (at least two consecutive) are expressed as percentages of its total in each group (see Table S2 for tallies).

### *Ms5788* is regulated in response to cysteine abundance

One predicted LL-sORF encodes eight consecutive cysteines at its 3’ end, standing out as a strong candidate for a role in cysteine attenuation (Table S2). We designate this polycysteine LL-sORF, *Ms5788A*, as it is immediately upstream of the annotated gene, *Ms5788*, which is followed by operonic genes, including a putative thiosulfate sulfurtransferase, *cysA2* (Fig 3)(Shell et al., 2015). RNA-seq and Ribo-seq profiles indicate that this unannotated LL-sORF is abundantly transcribed and translated in *M. smegmatis* (Fig S1A). Similar transcriptomic approaches in *M. tuberculosis* strongly suggest that a homologous LL-sORF, designated *Rv0815A*, is expressed in that species (Smith et al., 2019)(Fig S1B).

**Figure 3.**
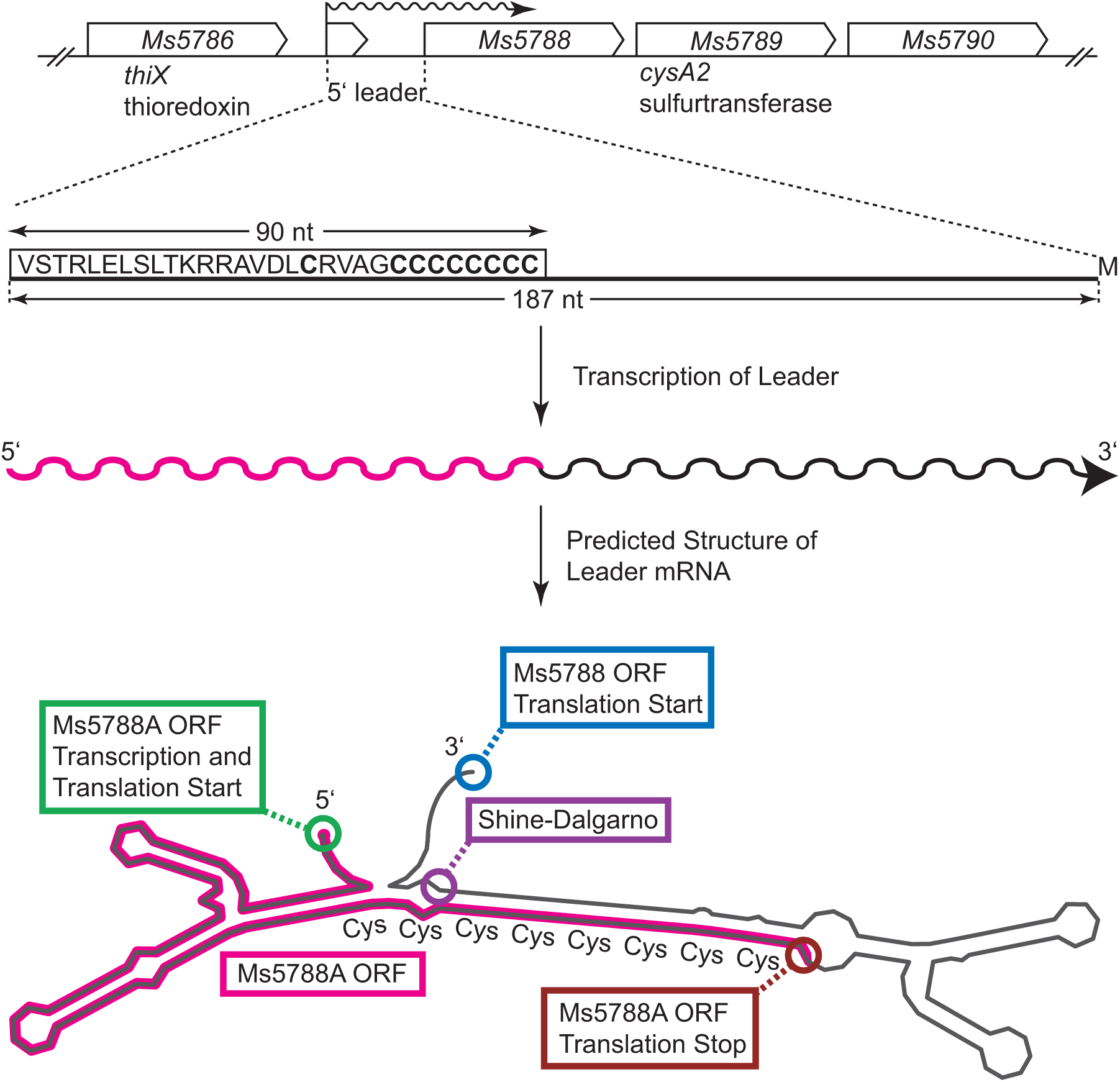
The architecture of the *Ms5788A* to *Ms5790* locus and the polycysteine-encoding mRNA leader. The amino acid sequence of the wild-type Ms5788A includes 9 cysteine codons (bold C), with eight consecutive codons at the C-terminus. This leader sequence to *Ms5788* is predicted to fold into a stable duplex structure (Fig S2), represented here in schematic form and annotated with features relevant for regulated expression of *Ms5788*. The ORF defined by the initiating GUG fMet is shown as a magenta line, with key sequences indicated.

The 90 nt *Ms5788A* LL-sORF and its 97 nt 3’ untranslated region (3’ UTR) comprise a 5’ leader of gene *Ms5788* (Fig 3). This RNA leader was analyzed by RNAfold to identify likely secondary structures. Nearly all of this compound leader sequence is predicted to form a stable duplex structure (Fig S2). In contrast to model attenuation systems that rely on competing terminator/anti-terminator stem loops of moderate stability, this predicted structure appears to be exceptionally stable. Moreover, the polycysteine tract encoded at the 3’ end of *Ms5788A* is predicted to base-pair with the region immediately preceding *Ms5788*, including the *Ms5788* Shine-Dalgarno (SD) sequence (Fig 3 and Fig S2). The predicted structure of the mRNA leader led us to hypothesize that under conditions of ample charged cysteine tRNA (tRNA^cys^), ribosomes translating Ms5788A quickly complete the process and release the mRNA at the stop codon (Fig 4A, right). As a consequence, the unoccupied cysteine tract of the sORF would be available for stable pairing with the SD region upstream of *Ms5788*. Under conditions of limiting cysteine, however, lower levels of charged tRNA^cys^ likely stall ribosomes in the polycysteine tract, preventing base-pairing and stable duplex formation (Fig 4A, left). Thus, ribosome-occupied sORF mRNA would not be available to base-pair with the *Ms5788* SD region, leaving the latter free to recruit ribosomes for translation of *Ms5788*.

**Figure 4.**
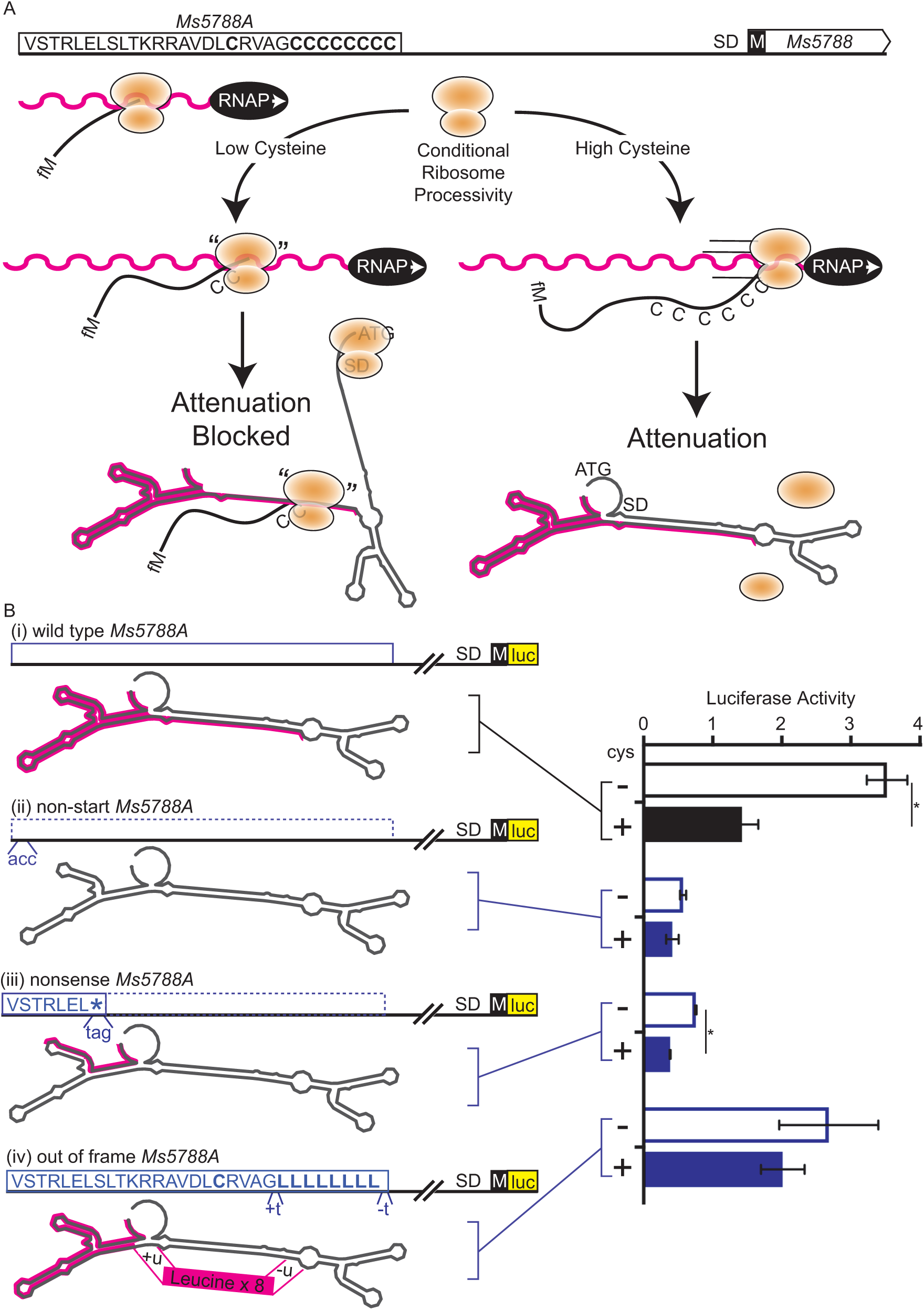
Hypothesized model and ribosome dependence of *Ms5788A*-directed attenuation. A. *Ms5788A* is transcribed (RNA polymerase, RNAP) and translated (orange ribosome). Ribosomes waiting (left) for charged tRNA^cys^ prevent the leader from folding into a stable, attenuating, conformation (right). The nascent small protein is shown emerging from the elongating ribosome and is annotated with the initiating methionine (fM) and polycysteine (C) tract for reference. Except where noted, luciferase reporters are translational fusions of the NanoLuc ORF in-frame with the AUG initiating methionine of the annotated *Ms5788* gene. B. The same wild-type (i) reporter cysteine responsive data are shown in Figs 4-6 (black outlined or filled boxes) as a comparative reference for the mutant derivatives of each series (blue outlined or filled boxes in Fig 4). Non-start (ii) and nonsense (iii) mutations reduce ribosome occupancy and increase attenuation of the reporter. An out-of-frame mutation permits ribosome occupancy but is not cysteine responsive (iv). Lower case letters below clones indicate nucleotide substitutions and insertion/deletions (+/-) used to create mutants. Luciferase activity is shown as light units (x 10^9^) per ml of cell culture, and asterisks indicate significance *p* <.01 for cysteine supplementation, by two-tailed t-test in Figs 4-6.

To test whether *Ms5788A* regulates *Ms5788* expression in response to changes in cysteine levels, we generated a luciferase translational reporter in which a constitutive promoter drives a leaderless transcript that begins at the native GUG initiation codon of *Ms5788A* and continues to the AUG initiation codon of *Ms5788* (Fig 4A, top). Minimal medium does not provide exogenous cysteine, requiring cells to activate the genes needed for its biosynthesis. We cultured *M. smegmatis* in minimal medium, with or without cysteine supplementation, and measured expression of the luciferase reporter. Luciferase activity decreased for cells grown with cysteine supplementation (Fig 4Bi). Hence, we hypothesized that expression of *Ms5788* is regulated in response to cysteine levels by an attenuation mechanism involving the upstream LL-sORF *Ms5788A* (Fig 1B).

We next tested whether cysteine-dependent regulation of *Ms5788* requires translation of the upstream *Ms5788A* LL-sORF. We mutated the GUG translation initiation codon of *Ms5788A* to ACC, to prevent ribosome loading. This non-start mutation reduced luciferase expression, and abolished attenuation (Fig 4Bii). These data indicate that translation of *Ms5788A* is required for cysteine responsiveness, and that in the absence of *Ms5788A* translation, expression of *Ms5788* is locked in an attenuated state. We also tested a nonsense mutant that truncated *Ms5788A* ten amino acids before the first Cys codon. This truncating mutation also reduced luciferase expression and attenuation by cysteine (Fig 4Biii). The residual cysteine response of this nonsense mutant may result from translation reinitiation after the stop codon. Thus, our data support a model in which cysteine-responsive regulation of the downstream gene requires ribosome occupancy of the LL-sORF.

We hypothesized that recoding the polycysteine tract would still allow ribosome occupancy but would no longer respond to cysteine. We created an out-of-frame (OoF) *Ms5788A* mutant reporter to recode the polycysteine tract to polyleucine. Supplementation with cysteine had no effect on this OoF reporter, which demonstrates that cysteine levels are monitored by the encoded *Ms5788A* polycysteine tract to modulate *Ms5788* gene expression (Fig 4Biv).

### Attenuation requires formation of a stable RNA duplex in the *Ms5788* 5’ UTR

The experiments above establish that ribosome occupancy of the polycysteine tract of *Ms5788A* relieves attenuated expression of the downstream reporter. Whereas the mutant series in Fig 4B were predicted to have minimal effect on mRNA structure, here we created mutants intended to selectively disrupt predicted duplex formation. In the first series (Fig 5, ii-iv), we changed the invariant guanine in the 2^nd^ position of cysteine codons (U<u>G</u>Y) to cytosine (U<u>C</u>Y) in the last five codons of *Ms5788A* (Fig 5ii). The nucleotide changes should reduce the stability of base-pair interactions with the SD region, while recoding polyserine for the final five codons of *Ms5788A*. This reporter exhibited constitutively elevated expression (Fig 5ii), consistent with unrestricted accessibility of the SD sequence regardless of tRNA^cys^ level.

**Figure 5.**
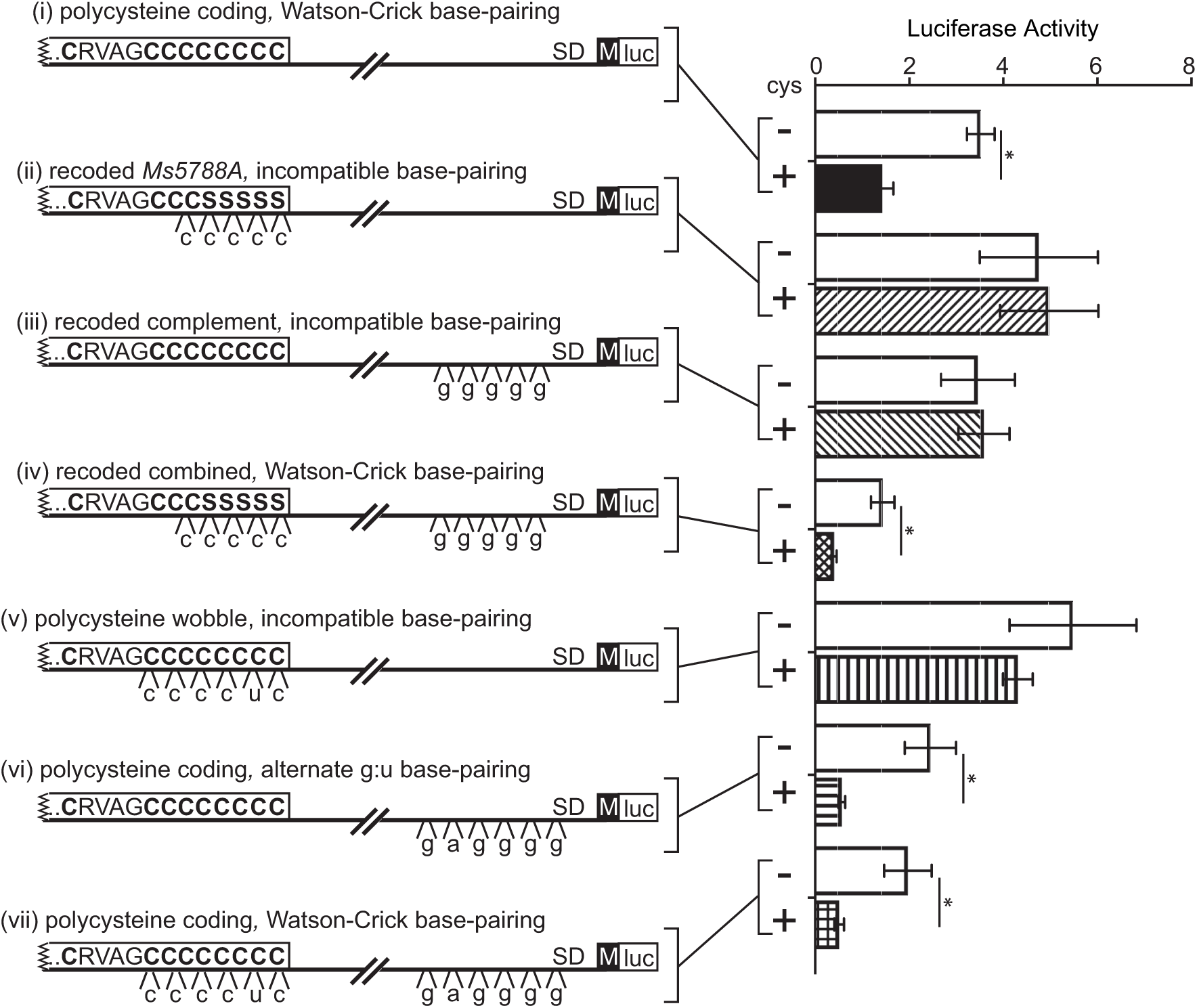
Predicted duplex mRNA interactions are required for the attenuated state of the *Ms5788* luciferase reporter fusion. Clustered point mutations were introduced into the *Ms5788A* LL-sORF or the predicted pairing nucleotides to disrupt base-pairing but, when combined, should restore the modeled stable mRNA structure. Two series of mutants were generated (angled hatch fill, ii – iv and horizontal and vertical hatch fill, v – vii). The effect of structure-destabilizing mutations and a re-stabilizing combination were tested for their effect on the luciferase reporter fused to *Ms5788.* All three clustered mutants (ii, iii, v) that are expected to disrupt the predicted duplex structure model an unattenuated state, whereas a G:U stabilized mutant (vi) or combined complement mutants (iv, vii) retain cysteine responsiveness. See Fig S3 for G:U stabilized structure detail.

To differentiate the effect of RNA duplex formation from the effect of *Ms5788A* amino acid recoding, we created a mutant that only disrupted the base-pairing by changing the predicted five cognate nucleotides near the SD sequence (Fig 5iii). Even though the polycysteine tract remained intact, luciferase expression from this reporter was insensitive to cysteine supplementation (Fig 5iii). The lack of cysteine response in this mutant is consistent with the nucleotide changes disrupting base-pairing, such that attenuation could not be achieved with excess cysteine. To restore base-pairing we next constructed a mutant that combined the recoded *Ms5788A* and cognate non-coding mutants. As expected, combining the complementary mutations reduced expression of the luciferase reporter for cells grown in the presence of cysteine, consistent with restored base-pairing (Fig 5iv). Interestingly, the restored response to cysteine suggests that the residual four-cysteine codon content of the LL-sORF is sufficient to confer sensitivity to cysteine.

In a second series (Fig 5, v-vii), we retained the full polycysteine tract and constructed a silent *Ms5788A* mutant that switched the codon choice of six cysteine codons to disrupt predicted base-pairing with the SD region. The elevated luciferase activity of this mutant reporter was not attenuated under cysteine-replete conditions, separating the roles of nucleotide and encoded amino acid sequence of *Ms5788A* (Fig 5v). We also created a mutant of the predicted complementary bases near the SD. Surprisingly, this mutant exhibited an attenuation response similar to wild type (Fig 5, compare i and vi). We reassessed the base-pairing potential of the mRNA produced from this mutant and found that new G:U base-pairings supported its folding into a stable, WT-like structure that would similarly reduce SD sequence availability under conditions of ample cysteine (Fig S3). Combining the *Ms5788A* and peri-SD nucleotide changes in this series resulted in an expression pattern similar to that of the WT construct, consistent with the model (Fig 5vii).

### Cysteine-dependent regulation involving *Ms5788A* affects expression of downstream operonic genes

Classical models of ribosome-mediated attenuation invoke competing mRNA stem loops that form an intrinsic terminator structure when ribosomes rapidly translate the sORF, resulting in regulation at the level of transcription (Turnbough, 2019, Yanofsky, 1981)(Fig 1). In the case of *Ms5788* attenuation, regulation occurs at the level of translation. To specifically test this, we created a reporter to assess mRNA transcription beyond the start of *Ms5788*. To maintain the predicted RNA structure of the 5’ leader region, an independent SD sequence was added to efficiently initiate translation of transcripts that extend beyond the modeled folded structure and into the luciferase gene (Fig 6). The added 15 nt sequence does not include identifiable complicating transcriptional or translational sequences. Luciferase activity of this reporter was insensitive to cysteine, and expression levels were consistently high (Fig 6ii). These data indicate that *Ms5788* attenuation does not result from transcription termination. Translational repression can indirectly affect transcription over longer distances due to polarity that is caused by Rho-dependent transcription termination and/or by enhanced RNase processing of the untranslated RNA (Deana and Belasco, 2005, Martini et al., 2019, Peters et al., 2011). To determine if translational repression of *Ms5788* by attenuation leads to polar effects on downstream genes, we constructed a translational fusion that extends to *Ms5789* (*cysA2*). This more distal site, 516 nt 3’ of the *Ms5788* start, exhibited cysteine responsiveness (Fig 6iii). Therefore, the ambient cysteine response imparted by *Ms5788A* to *Ms5788* translation initiation is also reflected in *Ms5789* gene expression. Taken together, our data support a model in which *Ms5788* expression is regulated by translational attenuation through *Ms5788A*-controlled SD availability, while *Ms5789* is likely regulated by polarity, due to the absence of elongating ribosomes in *Ms5788*.

**Figure 6.**
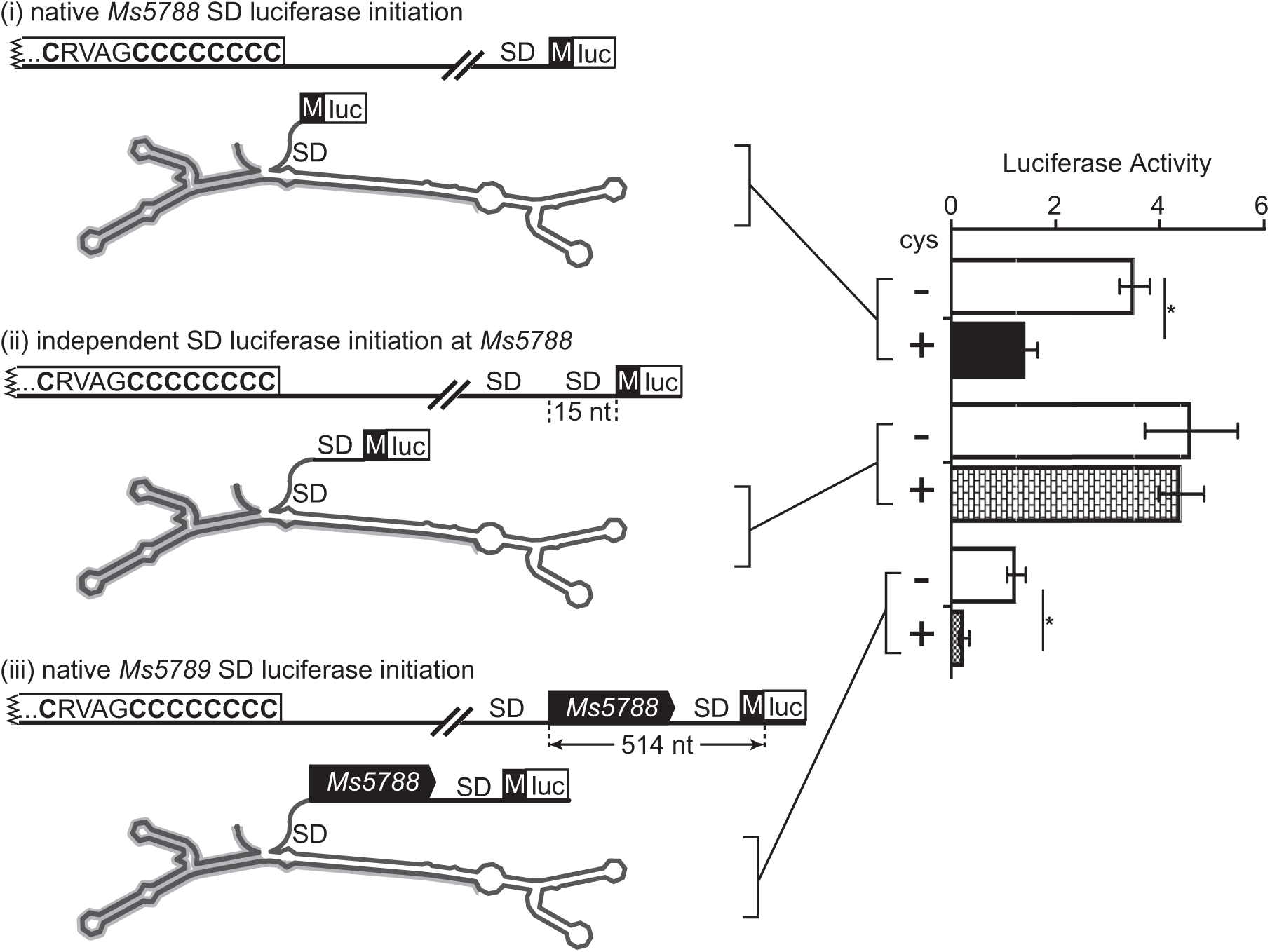
Attenuation regulates translation initiation of *Ms5788*. Fifteen nt were inserted at the end of the wild-type leader and included an independent SD to assess mRNA extension beyond the structured attenuator (ii). Luciferase activity is now independent of the effects of cysteine supplementation on the attenuating structure. Placement of the luciferase reporter at the initiation codon of *Ms5789* (iii) indicates that translation attenuation in *Ms5788* has polar effects, propagating the control of *Ms5788A* to operonic genes.

### Cysteine-dependent regulation of *Ms5789* and *Ms5790* by *Ms5788A* occurs in a chromosomal context

To assess whether our reporter-supported model is valid in the context of the native locus, we performed label-free quantitative (LFQ) mass spectrometry-based proteomic analysis to determine differences in protein expression for cells grown with or without cysteine supplementation. Whole cell extracts were prepared from cultures of *M. smegmatis* grown in minimal media +/- cysteine supplementation. Tryptic digests of whole cell lysates were subjected to nanoUHPLC-MS/MS. Peptides were identified and quantitated using LFQ based peak integration (Cox and Mann, 2008, Bosserman et al., 2017). As expected, the abundance of most proteins is unchanged between the two conditions (+/- cysteine), reflected in the linear correlation on the diagonal of the scatter plot (Fig 7A). Proteins below the diagonal were more abundant in cells grown without cysteine supplementation. The small (159 AA) and hydrophobic Ms5788 was not detected in these experiments. However, *Ms5789* and *Ms5790* are predicted to be co-transcribed in an operon with *Ms5788* (Martini et al., 2019), and expression of the encoded proteins Ms5789 and Ms5790 was higher in cells grown without cysteine (Fig 7A, diamonds 1 and 2), consistent with release from attenuation through *Ms5788A*.

**Figure 7.**
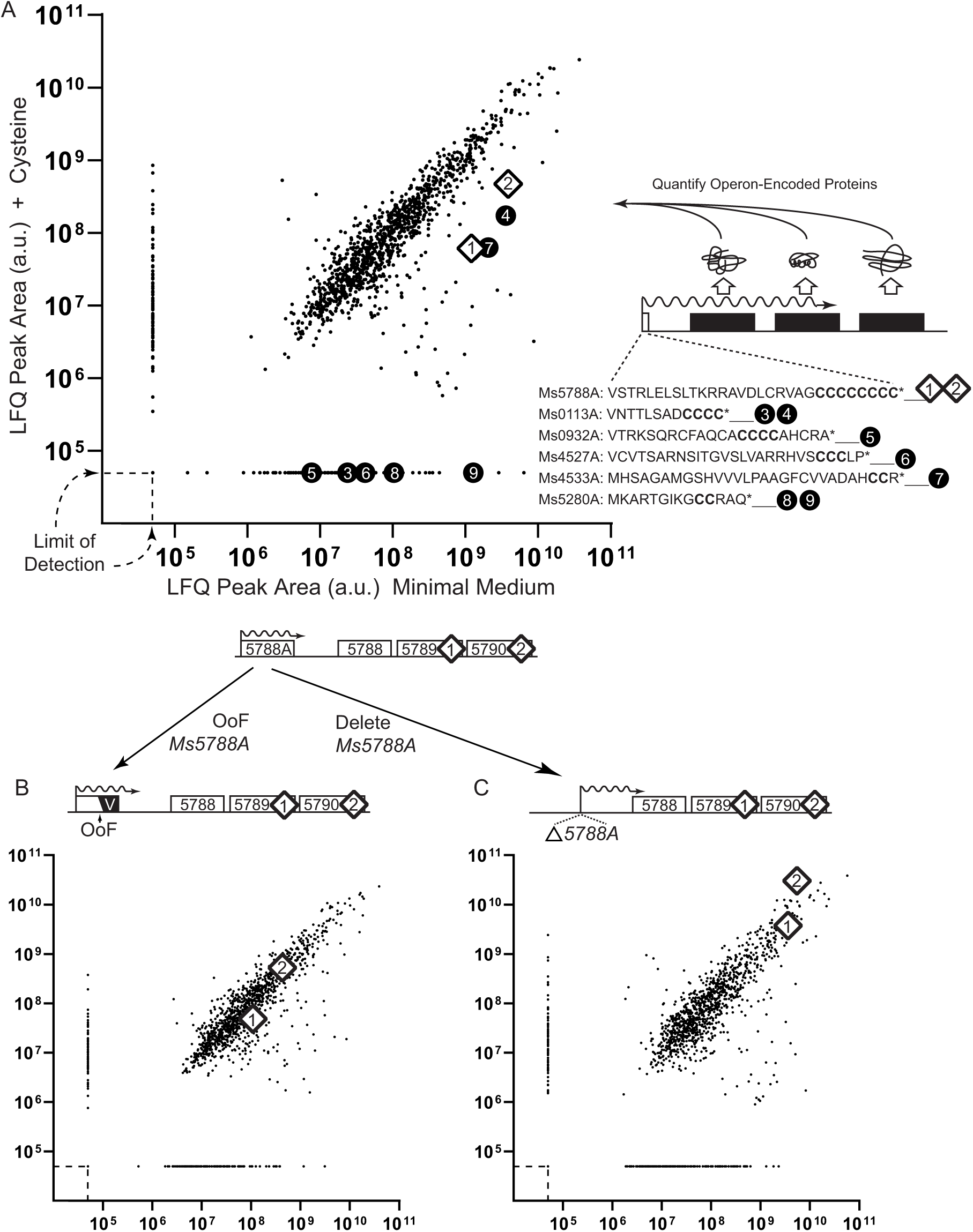
Cysteine attenuation is observed in the native context. A. Label-Free Quantitative proteomics (LFQ) was used to identify and quantitate changes in the abundance of proteins from whole-cell extracts of wild-type *M. smegmatis* subjected to trypsin digestion and nanoUHPLC-MS/MS. *M. smegmatis* was cultured in minimal medium (X-axis) or minimal medium supplemented with cysteine (Y-axis). Values plotted are normalized LFQ peak area (protein) in arbitrary units. The baselines for both X and Y-axes were artificially set at 5 ×10^4^ counts (indicated by dashed limit of detection) to allow depiction of proteins not expressed in one of the two conditions. Peptides from Ms5789 and Ms5790 (white diamond 1 and 2, respectively) increased in abundance (shift right along the X-axis) under cysteine limitation. Peptides of annotated proteins from loci similarly encoded on polycysteine-encoding LL-sORF mRNAs are indicated by black filled circles (Fig S4; 3 = Ms0113; 4 = Ms0114; 5 = Ms0934; 6 = Ms4527; 7 = Ms4533; 8 = Ms5279; 9 = Ms5280). B. A frameshift mutation (OoF) in *Ms5788A* that changes the CCCCCCCC* to VAVVVVAVERSRAL*. This OoF LL-sORF mutant is insensitive to cysteine and does not release Ms5789 and Ms5790 from an attenuated state. C. Deleting *Ms5788A* elevates expression of Ms5789 and Ms5790 and is insensitive to cysteine, indicating a completely unattenuated state.

To test the hypothesis that polycysteines in *Ms5788A* control Ms5789 and Ms5790 expression, we generated an out-of-frame (OoF) mutation in *Ms5788A* by deletion of a single nucleotide from the chromosomal locus. This deletion shifts the reading frame to encode valines and alanines in place of the polycysteine tract, and it adds six additional amino acids (ERSRAL) prior to encountering a stop codon (Fig 7B), while preserving the potential for nearly wild-type RNA duplex formation. This mutant exhibited lower Ms5789 and Ms5790 expression levels that were unaffected by cysteine supplementation (Fig 7B, diamonds move into the diagonal). This result links a point mutation in *Ms5788A* with gene expression of *Ms5789* and *Ms5790*, functionally demonstrating the polycistronic structure of this operon that is controlled by a LL-sORF.

Our working model suggests that if *Ms5788A* directs attenuation, its deletion would elevate operon expression of the downstream genes. We created a precise chromosomal deletion of *Ms5788A* and subjected this mutant to LFQ proteomics. As predicted, Ms5789 and Ms5790 were no longer responsive to ambient cysteine supplementation, appearing in the diagonal of unresponsive genes (Fig 7C, diamonds 1 and 2 now in the diagonal). Moreover, absolute levels of Ms5789 and Ms5790 were higher in the Δ*Ms5788A* strain than in either the wild-type or the OoF mutant, regardless of cysteine supplementation. This indicates that the *Ms5788A* deletion leaves the *Ms5788* SD sequence fully available for canonical translation initiation, and results in an elevated expression of the operonic Ms5789 and Ms5790 (Fig 7B, C diamonds 1 and 2). Thus, data from wild-type and targeted chromosomal mutant *M. smegmatis* are in full agreement with our reporter-generated data and strengthen the conclusion that the *Ms5788A* LL-sORF modulates operonic gene expression through a polycysteine-relieved attenuation mechanism.

### Widespread cysteine-dependent regulation in *M. smegmatis* associated with cysteine-rich LL-sORFs

While our in-depth analysis focused on the LL-sORF with the longest polycysteine tract, we hypothesized that other polycysteine LL-sORFs also regulate operonic gene expression in response to cysteine levels. We identified six more LL-sORFs that contain at least two consecutive cysteine codons and are located upstream of annotated genes. We then examined our LFQ proteomic data for the protein levels encoded by putative operonic genes downstream of these LL-sORFs for cells grown with/without cysteine supplementation. Remarkably, detectable proteins encoded by operons led by polycysteine LL-sORFs were also upregulated in low cysteine, falling below the diagonal (Fig 7A, black circles). Similar to *Ms5788A* controlling operonic expression of Ms5789 and Ms5790, these other LL-sORFs appear to control the expression of their operonic genes in response to cysteine supplementation. In cysteine-replete medium, expression of some of these proteins (Ms0113, Ms0934, Ms4527, Ms5279, and Ms5280) was below the threshold of detection for reliable quantification [<10^5^ LFQ intensity (a.u.)], indicative of tight attenuation. In the low-cysteine environment of minimal medium, attenuation at these polycysteine LL-sORF loci is apparently relieved, resulting in increased production of the operon-encoded proteins. The polycysteine LL-sORFs exhibit RNA-seq and Ribo-seq expression profiles consistent with their robust expression during cysteine replete growth (Figs S1 and S4). None of these expressed LL-sORFs were identified by genome annotation pipelines. Each of these cysteine-responsive operons contains genes annotated for cysteine associated activities (Table 1). Collectively, these data reveal a cysteine-metabolic regulon, whose concerted response is controlled independently at each locus by an expressed LL-sORF that includes consecutive cysteine codons.

**Table 1.**
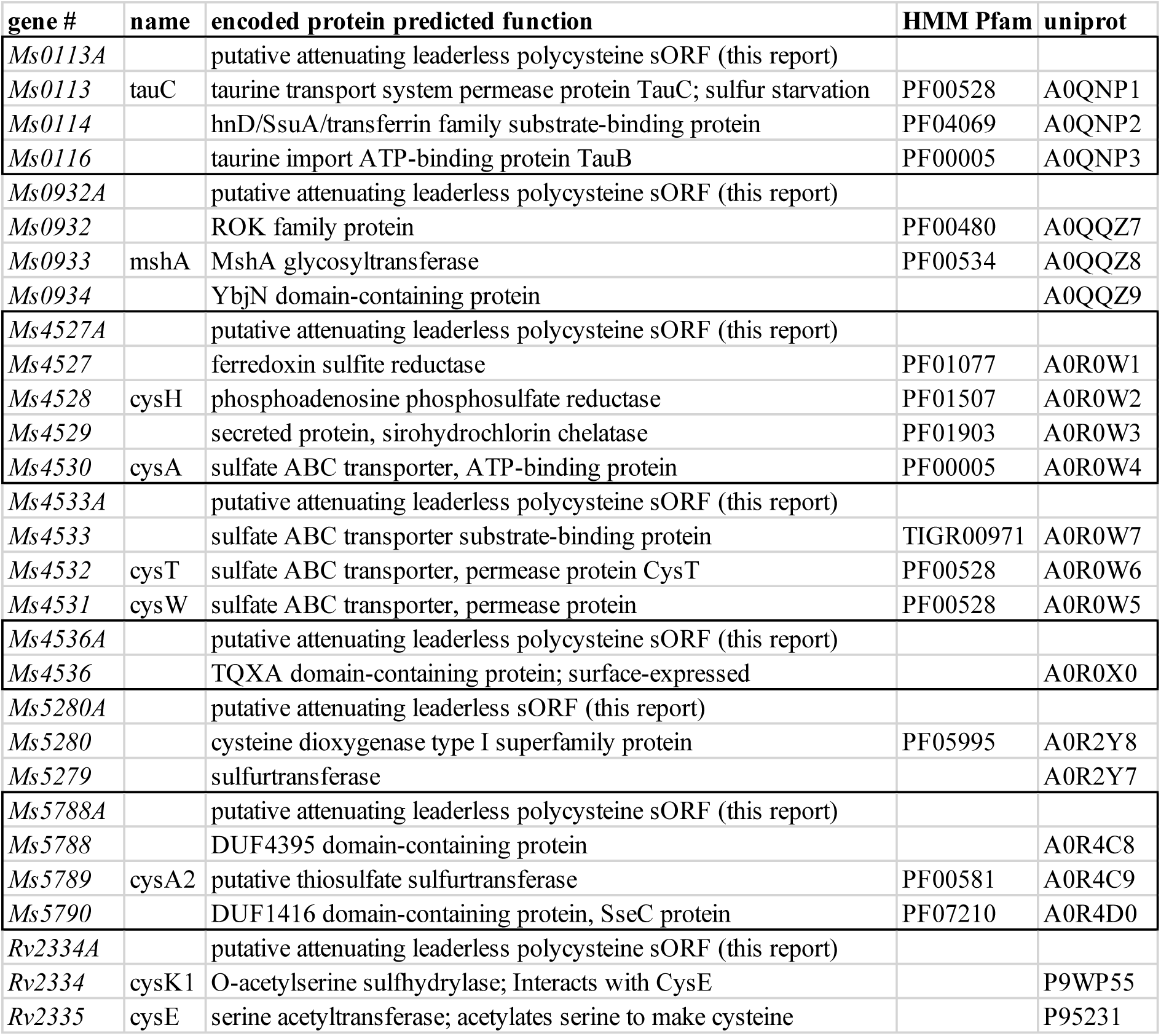
Cysteine translation attenuation regulon gene annotations. Gene order, content, and identifiers are listed each putative regulon operon for *M. smegmatis* (Ms). Only the *Rv2334A* operon is shown for *M. tuberculosis* (Rv). Operons are alternately boxed for clarity of separation. The predicted functions and features of the encoded proteins are shown (Pfam or uniprot).

LFQ proteomics does not comprehensively identify all proteins in a proteome. We looked for additional loci with the same hallmarks of responsive attenuation: an expressed polycysteine LL-sORF upstream of annotated operonic genes. *Ms4536A* is a transcribed polycysteine-encoding LL-sORF that was identified upstream of a single gene whose protein product, Ms4536/TQXA, was not detected by proteomics (Table 2, Tables S1 and S3, Fig S4E). Without corroborating indicators of cysteine response or function of the encoded operon protein, we can only speculate at its membership in the *M. smegmatis* cysteine LL-sORF regulon. The independent evolution of LL-sORFs makes it unlikely that our experimental reference species, *M. smegmatis*, contains all of the LL-sORF operons in the mycobacterial pan-genome. In one example, transcriptomic profile data for *M. tuberculosis* clearly identified *Rv2334A* as an unannotated gene meeting all of the criteria demonstrated in the *M. smegmatis* regulon operons: an expressed LL-sORF with a C-terminal polycysteine tract, followed by genes annotated as *cysK1* and *cysE* (Table 2, Fig S4G).

The availability of complete genome sequence information on diverse mycobacteria provided an opportunity to track the evolution of these polycysteine sORFs. We searched the genomes of 41 mycobacterial species using complementary approaches predicated on the sequence of each *M. smegmatis* or *M. tuberculosis* LL-sORF. The inherently low informatic content of sORFs make them poor BLAST queries, so we also used the position of the flanking annotated orthologous genes as landmarks. We identified genomic sequences consistent with conservation of the LL-sORFs expressed in *M. smegmatis* and *M. tuberculosis* (Table S3). The distribution of the conserved LL-sORFs is summarized as barcodes adjacent to each species on the *Mycobacterium* genus phylogenetic tree (Fig 8A). Sequence logos derived from multiple alignments of the amino acid sequences encoded by the LL-sORFs further support the evolutionary selection of their consecutive cysteine tracts (Fig 8B). The similarity between Ms4536A and Rv2334A strongly suggests homology by common origin, yet the context, including operon genes, is not homologous, suggesting origin by rearrangement rather than a merodiploid duplication or horizontal gene transfer event. Non-cysteine amino acids are also conserved in some LL-sORFs, suggesting that they provide function through ribosomal interaction, or support activities as an independent small protein product, or are encoded by codons that are constrained at the nucleotide level.

**Figure 8.**
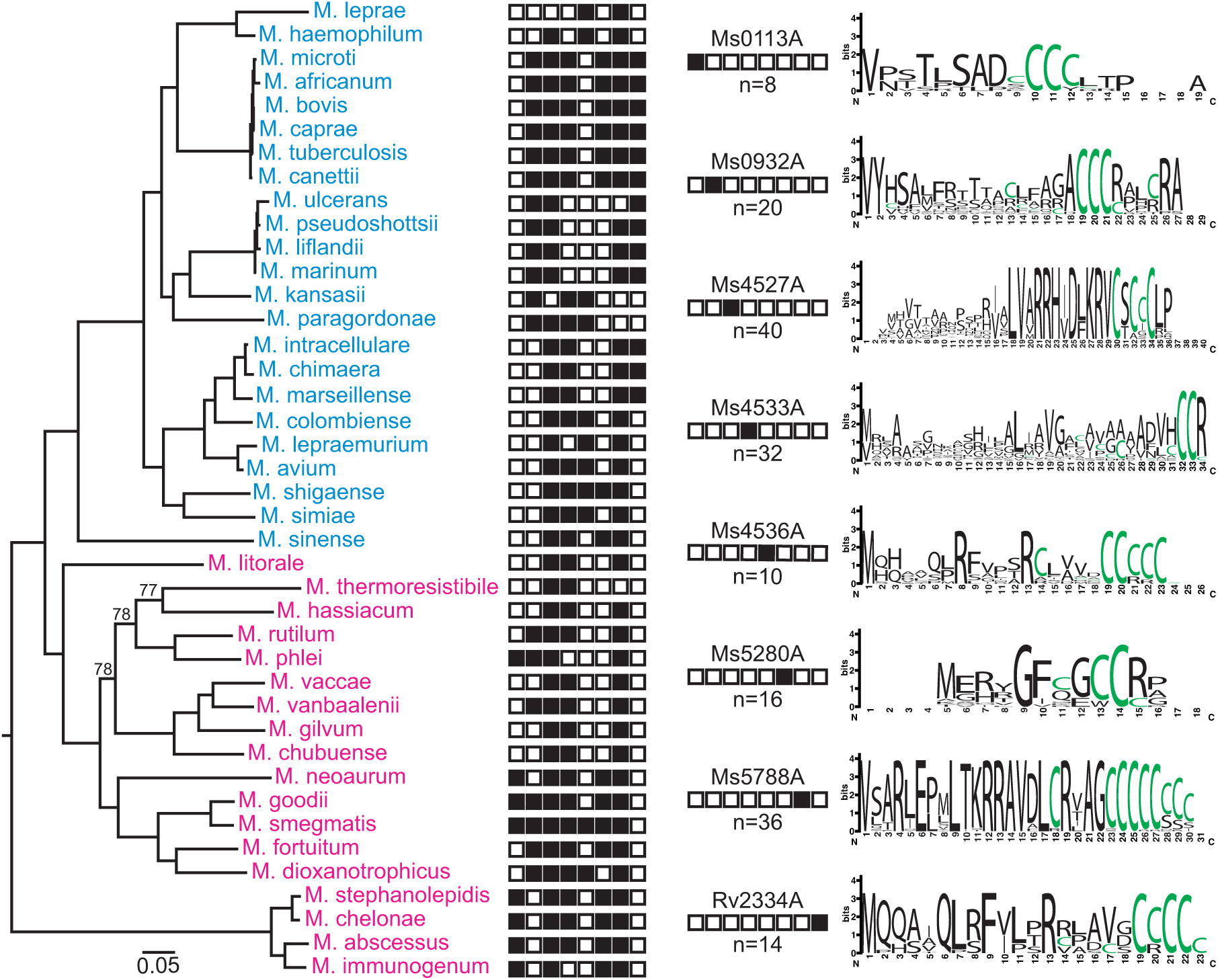
Phylogenetic distribution of polycysteine LL-sORFs in mycobacteria. A robust reference phylogenetic tree was constructed from complete genome sequences of 41 diverse *Mycobacterium* spp. All nodes had bootstrap values of 100, except where indicated. Species of the slow-grower clade appear in blue text, and fast-growers in magenta. The presence (black square) or absence (white square) of each LL-sORF forms a binary barcode for each species. For example, the presence of *Ms0113A* in some fast-growing species is indicated by a black square in the leftmost box of the barcode. Barcode key and fixed-length sequence logo for each LL-sORF is shown at right.

## DISCUSSION

### An attenuation mechanism that controls translation in response to amino acid availability

In the work presented here, we demonstrate attenuation of a cysteine biosynthesis pathway locus. Attenuation is a recurring theme in biosynthetic pathways for nucleosides and amino acids, in which the end product of the pathway interacts with the 5’ leader of the biosynthesis-encoding transcript to reduce (attenuate) expression of the operonic ORFs downstream (Turnbough, 2019). However, these models typically involve the formation of competing alternate hairpin structures that function as an intrinsic terminator when the end product is plentiful (Fig 1A). By contrast, the *Ms5788A* LL-sORF featured here defines a class of translational attenuator that indirectly assesses charged tRNA^cys^ availability to modulate expression of the downstream operonic genes. Blocking ribosome access to the SD sequence is a translational attenuation mechanism reported for sORF control of macrolide antibiotic resistance operons (Fig 1B) (Ito and Chiba, 2013). In the exit tunnel of the elongating ribosome, erythromycin interacts with the nascent peptide of the ermCL sORF of *Staphylococcus aureus*; the antibiotic-arrested ribosome alters mRNA hairpin structures to free the SD, allowing translation initiation of the *ermC* gene (Arenz et al., 2014). Chloramphenicol has a similar effect on the *cat* operon in *Bacillus subtilis* (Gu et al., 1994). These shared, yet varied, mechanisms illustrate how ribosome health is actively assessed and remedied, whether the cause is antibitioic attack or amino acid starvation.

### The need for cysteine regulation in mycobacteria

Why might mycobacteria need a multi-locus cysteine regulon? Mycobacteria rely on a cysteine derivative, mycothiol (MSH), to modulate redox balance (Loi et al., 2015, Xu et al., 2011). The *mshA* gene (*Ms0933*) encodes the first enzyme in MSH biosynthesis, and it resides in an operon with the hallmarks of cysteine attenuation (Fig 7A, Fig S4B). The multiple roles of cysteine in mycobacterial protein synthesis, redox and sulfur metabolism likely require subtle, independent fine-tuning of the enzymes involved in these respective pathways. The slight differences in cysteine composition and architectures of the polycysteine LL-sORFs could impart varying mechanisms as well as customized levels of operon gene expression.

### Attenuating sORF requirements

What are the cysteine codon requirements of an effective small ORF attenuator? Our criterion that LL-sORFs encode two consecutive cysteine codons may seem to be a low threshold, yet *cis*-encoded proteins only detected in unsupplemented minimal medium (baseline of Fig 7A) demonstrate the effectiveness of two consecutive cysteines in LL-sORFs in relieving attenuation. Each of the five LL-sORFs encoding at least three consecutive Cys codons (Table S2) were found to direct a cysteine response of their operonic proteins. Our *Ms5788A* LL-sORF analysis demonstrated that base-pairing was needed to impose attenuation, but it is premature to speculate that similar structures are required at all responsive loci. The additional cysteine-responsive operons that we identify are all led by an expressed polycysteine LL-sORF, but whether each attenuation mechanism is transcriptional or translational has not yet been determined. Regulation of transcription initiation in response to cysteine levels does not appear to be a factor at the *Ms5788* locus; the promoter remained intact in the *Ms5788A* mutants in Fig 7B, C, indicating that all of the cysteine response at this locus is directed by the attenuation mechanism we detailed and not via transcriptional activation. RNA-seq profiles of the LL-sORFs presented here (Fig S1, S4) are also consistent with robust constitutive transcription in the cysteine-replete medium used for routine culturing.

### Evolution of a cysteine attenuation regulon; the Cys ribulon

The independent evolution of similar polycysteine LL-sORF architecture associated with cysteine attenuation at multiple loci in mycobacteria indicates that coordinated expression is important, and that LL-sORF-directed attenuation is effective. The co-regulation of individual operons, caused by ribosome stalling at sORFs that are cysteine sensors in this scenario, functionally defines a regulon that we term the “ribulon” (Fig 9). The term ribulon reflects that control is dictated by the ribosome pausing and not a protein *per se*, and it would be functionally analogous to regulons controlled by dedicated DNA-binding transcription factors. We speculate that the evolution of regulation by attenuation is simplified in mycobacteria by the robust nature of leaderless translation, and that the evolution of a dedicated transcription factor and its cognate binding sites in the promoters of target operons is more problematic than exploiting LL-sORFs in a genus that exhibits frequent and robust LL-mRNA expression. As the first genes in their transcripts, LL-mRNAs are ideally positioned to *cis*-regulate expression of downstream genes. Additionally, transcription start sites are preferentially associated with purines (R), and the +2 transcript position is preferentially associated with pyrimidines (Y) (Martini et al., 2019). Thus, transcription often begins at RYN trinucleotide sequences. Our previous study showed that a 5’ RUG trinucleotide confers robust leaderless translation initiation (Shell et al., 2015), so many transcription start sites are only one or two changes from also initiating translation. It is not yet clear whether leaderless architecture *per se* offers advantages for attenuation and may have been selected over canonical Shine-Dalgarno translation initiation, or whether the sole criterion of an RUG sequence at the transcription start site is simply a relatively modest requirement. It is clear, however, that in mycobacteria, polycysteine LL-sORFs integrate two levels of coordination: locally by modulating polycistronic operon gene expression, and globally by coordinating the response of cysteine-related operons.

**Figure 9.**
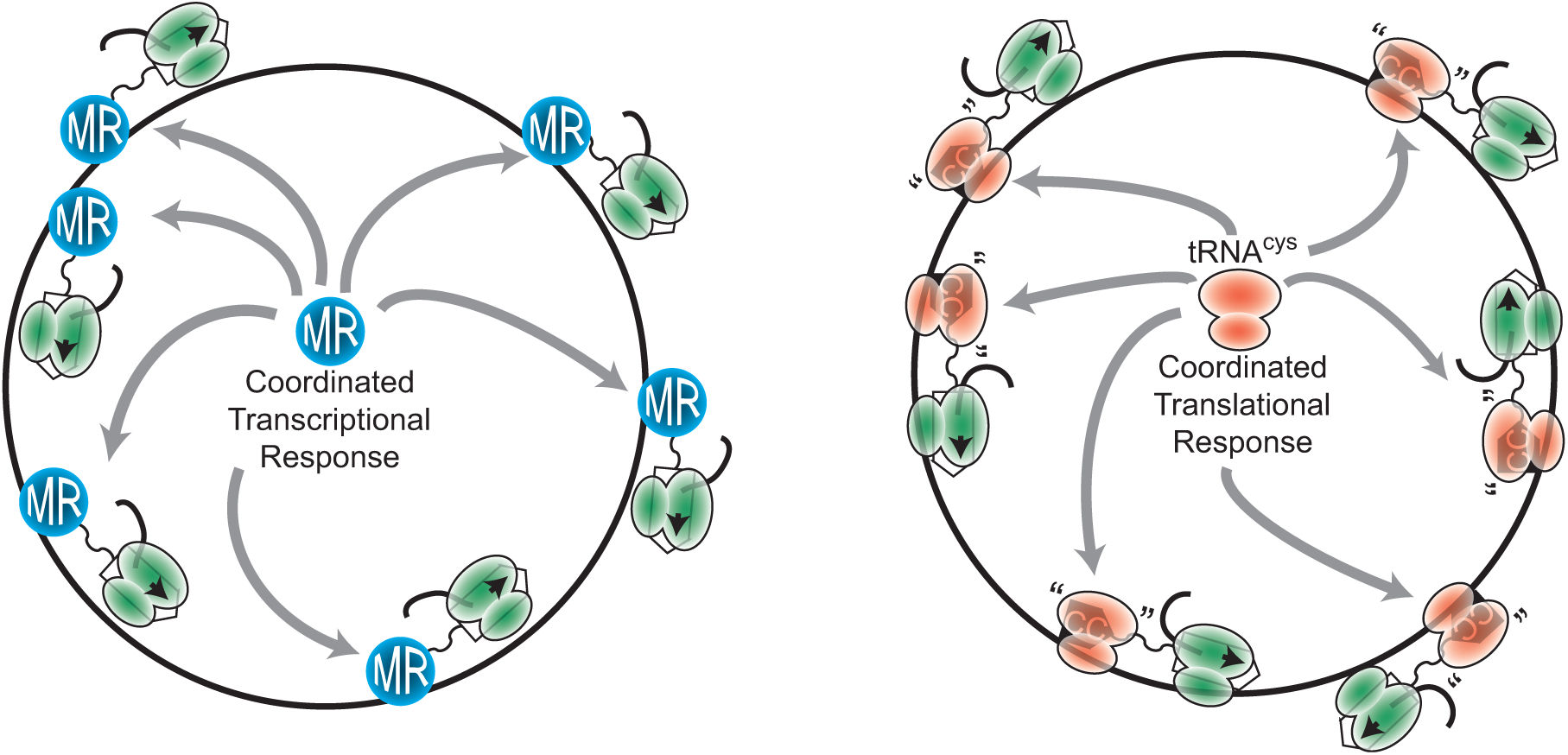
Schematic comparison of a traditional transcriptionally coordinated regulon and the translationally controlled ribulon. A. Regulons are usually controlled at the transcriptional level and are named for their master regulator (MR). In this scenario, coordinate gene expression from multiple loci, is mediated by the MR activating transcription. The MR is shown binding to multiple loci, which activates transcription and subsequent translation by the ribosome. This includes activation of sigma factors (e.g., Sigma F flagellar regulon in *Salmonella typhimurium* (Ohnishi et al., 1992)), inactivation of a repressor (e.g., *<u>fur</u>* regulon in *E. coli* (Stojiljkovic et al., 1994)), or two-component activation (e.g., *slyA* regulon in *Salmonella typhimurium* (Zhao et al., 2008)). B. In the Cys ribulon, global gene coordination occurs by mutual pausing of ribosomes in ribulon sORFs. Attenuation at individual LL-sORFs at independent operons is co-dependent on tRNA^cys^ levels, resulting in coordinated gene expression. In cysteine replete conditions, polycysteine sORFs are rapidly translated and expression of downstream genes is repressed. Under conditions of limiting tRNA^cys^ levels, ribosomes pause at polycysteine tracts in the sensory LL-sORFs and relieve attenuation of operonic genes to upregulate processes required for the production of cysteine. Sensory sORFs are shown as black boxes, stalled ribosomes are red-shaded, activated genes are indicated by open rectangles and translated by elongating ribosomes (green-shaded) with the emerging nascent protein.

### Concluding remarks

Given the prevalence of LL-sORFs in mycobacteria, we speculate that other translational regulons have evolved as an alternative to transcriptional regulons controlled by DNA-binding transcription factors. LL translation is considered to be the ancestral form of ribosome delivery (Duval et al., 2013, Nakamoto, 2009, Zheng et al., 2011), indicating that LL-sORF regulons may be ancient and widespread. We propose that LL-sORFs define a functional subclass of small *de novo* genes that effectively decode translational stress into broad regulatory effects.

## EXERIMENTAL PROCEDURES

### Bacterial strains and culture

*M. smegmatis* wild-type mc^2^155 and its derivatives were grown in tryptic soy broth + 0.05% Tween 80 (TSBT) or on TSA plates, and cultured at 37 °C. Antibiotic selection for reporter maintenance or mutation selection strategies included apramycin (12.5 µg ml^-1^ on agar, 10 µg ml^-1^ in broth), hygromycin (100 and 25 µg ml^-1^), kanamycin (50 and 10 µg ml^-1^), and zeocin (50 and 25 µg ml^-1^).

For the cysteine attenuation study, bacteria were cultured in minimal media. Base medium per liter: 6 g Na_2_HPO_4_ (anhydrous), 3 g KH_2_PO_4_, 0.5 g NaCl, 1 g NH_4_Cl, 0.05% (v/v) Tween-80. After autoclaving, 0.2 % glucose and micronutrients were added to final concentrations: MgSO_4_ to 1 mM, CaCl_2_ to 100 μM, H_3_BO_3_ to 4×10^−7^ M, CoCl_2_.6H_2_O to 3×10^−8^ M, CuSO_4_.5H_2_O to 1×10^−8^ M, MnCl_2_.4H_2_O to 8×10^−8^ M, ZnSO_4_.7H_2_O to 1×10^−8^ M, FeSO_4_.7H_2_O to 1×10^−6^ M. L-cysteine or L-cystine (exogenously stable dimeric cysteine) was supplemented at 200 µg ml^-1^ as noted.

### Luciferase assays

Reporters were generated by long-primer-dimer PCR to recreate the LL-sORF and leader sequence of *M. smegmatis Ms5788*. PCR products were cloned by Infusion (Clontech) and verified by DNA sequence analysis. The NanoLuc (Promega) luciferase ORF is carried on a plasmid that confers apramycin resistance and integrates at the L5 *attB* site in mycobacteria. Mycobacterial cultures were grown in minimal medium for luciferase assays. Luciferase activity in a culture was assessed by the addition of NanoGlo (Promega) substrate and then measuring luminescence normalized to culture density.

### Chromosomal mutants of *M. smegmatis*

A precise deletion of *Ms5788A* was created using a targeting plasmid that integrates via a single cross-over, allowing selection of a hyg^r^ intermediate and, then, is resolved by a second homology-driven event in which a *sacB*/*galK* counter-selection allows enrichment for the deleted recombinant (Barkan et al., 2011). The OoF point mutant was created by a recombineering approach that used a single-stranded oligonucleotide template to introduce an additional adenine in *Ms5788A* (van Kessel and Hatfull, 2008). Co-electroporation of 2 × 10^−10^ mol of 60-mer oligo with 500 ng of an episomal zeo^r^ plasmid (pGE324, Zeo-*sacB*) allowed zeocin selection and isolation of the electrocompetent population of *M. smegmatis*. Mismatch-sensitized PCR (MAMA-PCR) screening (Cha et al., 1992, Swaminathan et al., 2001) of isolates identified clones that integrated the additional adenine. Deletion and point mutants were verified by genomic DNA PCR and sequencing.

### Mass spectrometry

Wild-type and mutant derivative *M. smegmatis* were cultured in minimal media with or without cysteine supplementation, harvested by centrifugation and cryo-milled (Retsch MM400, Haan, Germany) for mass spectrometry analysis. Reagents were of LC-MS quality or higher and obtained from Sigma Aldrich unless indicated. Milled cell pellets were digested with trypsin using commercial S-Traps (Protifi, NY). Briefly 50 μg of protein from each milled cell pellet was re-suspended in 6% SDS 10 mM Tris-2-Carboxy ethyl phosphine in 100 mM tri ethyl ammonium bicarbonate (TEAB), heated at 95 °C for 3 minutes, then alkylated with 10 mM iodoacetamide in the dark for 15 min. Samples were acidified by addition of H_3_PO4_4_ to 1.2% final concentration (v/v) and flocculated by 7-fold addition of 95:5 MeOH:100 mM TEAB prior to collection on an S-trap column (Zougman et al., 2014). Washing and conditioning was preformed three times by addition of 150 μ MeOH buffer, as described above. 1 μg of sequencing-grade trypsin (Promega, WI) was added to each sample and digested at 37 °C for 8 hours. Peptides were isolated, acidified and desalted using Stage tips packed into P100 pipette tips, and dried using a MiVac (Genvac UK) prior to LC-MS/MS analysis (Rappsilber et al., 2003).

NanoUHPLC-MS/MS was performed essentially as described (Bosserman et al., 2017, Bosserman et al., 2019). 1 µg of each digest was analyzed in technical triplicate and biological duplicate on an Orbitrap instrument running a TOP15 data-dependent acquisition (Q-Exactive Thermo San Jose, CA). Protein spectral matching and Label Free quantification were performed using MaxQuant (Cox and Mann, 2008) against the *M. smegmatis* FASTA combined with contaminants from the Uniprot database, LFQ param (UP000000757 6,595 entries). LFQ parameters were set to default, quantification was restricted to proteins >2 peptides (missing peaks enabled). Target-decoy was used to determine False Discovery Rates (Elias and Gygi, 2007) and proteins at a global 1% FDR were used for quantification. Data reduction and significance testing were performed using a modified LIMMA methodology (Efstathiou et al., 2017). Protein search and RAW data files are accessible at the Center for Computational Mass Spectrometry via ftp://MSV000084381@massive.ucsd.edu (Deutsch et al., 2017). Proteins quantitatively replicated in one condition but absent in another, were given an arbitrary intensity of 5×10^4^ (below detection threshold) for ease in visualization.

### Phylogenetic analysis

The genomes and proteomes for *Mycobacterium abscessus* subsp. massiliense (NC_018150.2), *M. africanum* strain 25 (CP010334,1), *M. litorale* strain F4 (CP019882.1), *M. avium* 104 (NC008595.1), *M. ulcerans* Agy99 (NC_008611.1), *M. vanbaalenii* PYR-1 (NC_008726.1), *M. marinum* M (NC_010612.1), *M. liflanddii* 128FXT (NC_020133.1), *M. kansasii* ATCC 12478 (NC_022663.1), *M. gilvum* Spyr1 (NC_014814.1), *M. bovis* AF2122/97 (NC_002945.4), *M. tuberculosis* H37Rv (NC_000963.3), *M. sinense* strain JDM601 (NC_015576.1), *M. canettii* CIPT 140010059 (NC_015848.1), *M. chubuense* NBB4 (NC_018027.1), *M. intracellulare* MOTT-64 (NC_016948.1), *M. intracellulare* ATCC 13950 (NC_016946.1), *M. neoaurum* VKM ac-1815D (NC_023036.2), *M. haemophilum* ATCC 29548 (NZ_CP0118883.2), *M. simiae* ATCC 25275 (NZ_HG315953.1), *M. goodii* strain X7B (NZ_CP012150.1), *M. fortuitum* strain CT6 (NZ_CP011269.1), *M. phlei* strain CCUG 21000 (NZ_CP014475.1), *M. immunogenum* strain CCUG 47286 (NZ_CP011530.1), *M. chelonae* CCUG 47445 (NZ_CP007220.1), *M. vaccae* 95051 (NZ_CP011491.1), *M. chimaera* strain AH16 (NZ_CP012885.2), *M. caprae* strain Allgaeu (NZ_CP016401.1), *M. colombiense* CECT 3035 (NZ_CP020821.1), *M. dioxanotrophicus* strain PH-06 (NZ_CP020809.1), *M. marseillense* strain FLAC0026 (NZ_CP023147.1), *M. lepraemurium* strain Hawaii (NZ_CP021238.1), *M. shigaense* strain UN-152 (NZ_AP018164.1), *M. stephanolepidis* (NZ_AP018165.1), *M. pseudoshottsii* JCM 15466 (NZ_AP018410.1), *M. paragordonae* 49061 (NZ_CP025546.1), *M. rutilum* strain DSM 45405 (NZ_LT629971.1), *M. thermoresistibile* strain NCTC10409 (NZ_LT906483.1), *M. hassiacum* DSM 44199 (NZ_LR026975.1), and *M. microti* strain 12 (CP010333.1) were downloaded from NCBI. Orthologs of *M. smegmatis* proteins were identified by a reciprocal best BLAST hit approach and aligned in MAFFT v.7.058b (Katoh and Standley 2013). A supermatrix was created from the concatenation of 1560 of these protein alignments with an in-house Python script. Alignments included in the supermatrix were required to have 40-41 of the 41 mycobacterial species present. IQ-TREE v.1.6.9 (Nguyen et al., 2015) generated a maximum likelihood (ML) phylogeny under an LG+R4 model of evolution (Soubrier et al., 2012; Yang 1995; Le and Gascuel 2008). Support values were generated by the ultrafast bootstrap method with 1000 replicates (Minh et al., 2013). The ML tree was visualized in FigTree v.1.4.4 (http://tree.bio.ed.ac.uk/software/figtree/).

### Identification of sORFs in additional mycobacterial genomes

The orthologs to all *M. smegmatis* ORFs with an upstream sORF were identified in other mycobacteria via a reciprocal best blast hit approach and the regions 1000 nucleotides upstream of these genes were extracted. *M. smegmatis* sORF proteins were queried against these upstream regions via tBLASTn searches to identify orthologous sequences. The coordinates for these sequences were obtained from the blast results and extended to the match the length of the corresponding *M. smegmatis* sORF. To ensure that we captured the start and stop codon, we extended these coordinates by an additional 5-10 amino acids on both 5’ and 3’ ends. These coordinates and the strand information captured from the blast results were used to extract and translate sORFs from 1 Kb upstream regions with an in-house Python script. All sORFs were visually inspected and initially aligned in MEGA v. 70.26 (Kumar et al., 2016). In some cases, additional sORF orthologs were identified by predicting proteins (anything between two stop codons) in upstream 1 Kb regions with the getorf subroutine from EMBOSS v. 6.3.1 (Rice et al., 2000).

Multiple alignments of deduced LL-sORF amino acid sequences were performed using a web-based tool and default clustal algorithm (Madeira et al., 2019). The aligned sequences were compiled into a sequence logo infographic (weblogo.berkeley.edu).

## Acknowledgments

This work was supported by NIH grants R21AI147608, R21AI119427, R01AI097191, R21AI117158 to KMD, TAG, JTW, and MMC, and by Wadsworth Center Cores for Bioinformatics & Statistics, Applied Genomic Technologies, and Media & Tissue Culture. We thank Dr. Paul Masters for his thoughtful comments in manuscript preparation.

## REFERENCES

Arenz, S., Meydan, S., Starosta, A. L., Berninghausen, O., Beckmann, R., Vazquez-Laslop, N. & Wilson, D. N. 2014. Drug sensing by the ribosome induces translational arrest via active site perturbation. Mol Cell, 56, 446–52.

Barkan, D., Stallings, C. L. & Glickman, M. S. 2011. An improved counterselectable marker system for mycobacterial recombination using galK and 2-deoxy-galactose. Gene, 470, 31–6.

Bechhofer, D. H. 1990. Triple post-transcriptional control. Mol Microbiol, 4, 1419–23.

Beck, H. J. & Moll, I. 2018. Leaderless mRNAs in the Spotlight: Ancient but Not Outdated! Microbiol Spectr, 6.

Bosserman, R. E., Nguyen, T. T., Sanchez, K. G., Chirakos, A. E., Ferrell, M. J., Thompson, C. R., Champion, M. M., Abramovitch, R. B. & Champion, P. A. 2017. WhiB6 regulation of ESX-1 gene expression is controlled by a negative feedback loop in Mycobacterium marinum. Proc Natl Acad Sci U S A, 114, E10772–E10781.

Bosserman, R. E., Nicholson, K. R., Champion, M. M. & Champion, P. A. 2019. A New ESX-1 Substrate in Mycobacterium marinum That Is Required for Hemolysis but Not Host Cell Lysis. J Bacteriol, 201.

Burian, J. & Thompson, C. J. 2018. Regulatory genes coordinating antibiotic-induced changes in promoter activity and early transcriptional termination of the mycobacterial intrinsic resistance gene whiB7. Mol Microbiol, 107, 402–415.

Cha, R. S., Zarbl, H., Keohavong, P. & Thilly, W. G. 1992. Mismatch amplification mutation assay (MAMA): application to the c-H-ras gene. PCR Methods Appl, 2, 14–20.

Cortes, T., Schubert, O. T., Rose, G., Arnvig, K. B., Comas, I., Aebersold, R. & Young, D. B. 2013. Genome-wide mapping of transcriptional start sites defines an extensive leaderless transcriptome in Mycobacterium tuberculosis. Cell reports, 5, 1121–31.

Couso, J. P. & Patraquim, P. 2017. Classification and function of small open reading frames. Nat Rev Mol Cell Biol, 18, 575–589.

Cox, J. & Mann, M. 2008. MaxQuant enables high peptide identification rates, individualized p.p.b.-range mass accuracies and proteome-wide protein quantification. Nat Biotechnol, 26, 1367–72.

Crappe, J., Van Criekinge, W., Trooskens, G., Hayakawa, E., Luyten, W., Baggerman, G. & Menschaert, G. 2013. Combining in silico prediction and ribosome profiling in a genome-wide search for novel putatively coding sORFs. BMC Genomics, 14, 648.

Deana, A. & Belasco, J. G. 2005. Lost in translation: the influence of ribosomes on bacterial mRNA decay. Genes Dev, 19, 2526–33.

Deutsch, E. W., Csordas, A., Sun, Z., Jarnuczak, A., Perez-Riverol, Y., Ternent, T., Campbell, D. S., Bernal-Llinares, M., Okuda, S., Kawano, S., Moritz, R. L., Carver, J. J., Wang, M., Ishihama, Y., Bandeira, N., Hermjakob, H. & Vizcaino, J. A. 2017. The ProteomeXchange consortium in 2017: supporting the cultural change in proteomics public data deposition. Nucleic Acids Res, 45, D1100–D1106.

Duval, M., Korepanov, A., Fuchsbauer, O., Fechter, P., Haller, A., Fabbretti, A., Choulier, L., Micura, R., Klaholz, B. P., Romby, P., Springer, M. & Marzi, S. 2013. Escherichia coli ribosomal protein S1 unfolds structured mRNAs onto the ribosome for active translation initiation. PLoS Biol, 11, e1001731.

Efstathiou, G., Antonakis, A. N., Pavlopoulos, G. A., Theodosiou, T., Divanach, P., Trudgian, D. C., Thomas, B., Papanikolaou, N., Aivaliotis, M., Acuto, O. & Iliopoulos, I. 2017. ProteoSign: an end-user online differential proteomics statistical analysis platform. Nucleic Acids Res, 45, W300–W306.

Elias, J. E. & Gygi, S. P. 2007. Target-decoy search strategy for increased confidence in large-scale protein identifications by mass spectrometry. Nat Methods, 4, 207–14.

Frith, M. C., Forrest, A. R., Nourbakhsh, E., Pang, K. C., Kai, C., Kawai, J., Carninci, P., Hayashizaki, Y., Bailey, T. L. & Grimmond, S. M. 2006. The abundance of short proteins in the mammalian proteome. PLoS Genet, 2, e52.

Gu, Z., Harrod, R., Rogers, E. J. & Lovett, P. S. 1994. Anti-peptidyl transferase leader peptides of attenuation-regulated chloramphenicol-resistance genes. Proc Natl Acad Sci U S A, 91, 5612–6.

Hemm, M. R., Paul, B. J., Miranda-Rios, J., Zhang, A., Soltanzad, N. & Storz, G. 2010. Small stress response proteins in Escherichia coli: proteins missed by classical proteomic studies. J Bacteriol, 192, 46–58.

Hemm, M. R., Paul, B. J., Schneider, T. D., Storz, G. & Rudd, K. E. 2008. Small membrane proteins found by comparative genomics and ribosome binding site models. Mol Microbiol, 70, 1487–501.

Henkin, T. M. & Yanofsky, C. 2002. Regulation by transcription attenuation in bacteria: how RNA provides instructions for transcription termination/antitermination decisions. Bioessays, 24, 700–7.

Hinnebusch, A. G., Ivanov, I. P. & Sonenberg, N. 2016. Translational control by 5’-untranslated regions of eukaryotic mRNAs. Science, 352, 1413–6.

Hobbs, E. C., Fontaine, F., Yin, X. & Storz, G. 2011. An expanding universe of small proteins. Current opinion in microbiology, 14, 167–73.

Ito, K. & Chiba, S. 2013. Arrest peptides: cis-acting modulators of translation. Annu Rev Biochem, 82, 171–202.

Johnston, H. M., Barnes, W. M., Chumley, F. G., Bossi, L. & Roth, J. R. 1980. Model for regulation of the histidine operon of Salmonella. Proc Natl Acad Sci U S A, 77, 508–12.

Loi, V. V., Rossius, M. & Antelmann, H. 2015. Redox regulation by reversible protein S-thiolation in bacteria. Front Microbiol, 6, 187.

Lovett, P. S. & Rogers, E. J. 1996. Ribosome regulation by the nascent peptide. Microbiological reviews, 60, 366–85.

Madeira, F., Park, Y. M., Lee, J., Buso, N., Gur, T., Madhusoodanan, N., Basutkar, P., Tivey, A. R. N., Potter, S. C., Finn, R. D. & Lopez, R. 2019. The EMBL-EBI search and sequence analysis tools APIs in 2019. Nucleic Acids Res, 47, W636–W641.

Martini, M. C., Zhou, Y., Sun, H. & Shell, S. S. 2019. Defining the Transcriptional and Post-transcriptional Landscapes of Mycobacterium smegmatis in Aerobic Growth and Hypoxia. Front Microbiol, 10, 591.

Meydan, S., Marks, J., Klepacki, D., Sharma, V., Baranov, P. V., Firth, A. E., Margus, T., Kefi, A., Vazquez-Laslop, N. & Mankin, A. S. 2019. Retapamulin-Assisted Ribosome Profiling Reveals the Alternative Bacterial Proteome. Mol Cell, 74, 481–493 e6.

Miranda-Casoluengo, A. A., Staunton, P. M., Dinan, A. M., Lohan, A. J. & Loftus, B. J. 2016. Functional characterization of the Mycobacterium abscessus genome coupled with condition specific transcriptomics reveals conserved molecular strategies for host adaptation and persistence. BMC Genomics, 17, 553.

Nakamoto, T. 2009. Evolution and the universality of the mechanism of initiation of protein synthesis. Gene, 432, 1–6.

Ohnishi, K., Kutsukake, K., Suzuki, H. & Lino, T. 1992. A novel transcriptional regulation mechanism in the flagellar regulon of Salmonella typhimurium: an antisigma factor inhibits the activity of the flagellum-specific sigma factor, sigma F. Mol Microbiol, 6, 3149–57.

Oppenheim, D. S. & Yanofsky, C. 1980. Translational coupling during expression of the tryptophan operon of Escherichia coli. Genetics, 95, 785–95.

Peters, J. M., Vangeloff, A. D. & Landick, R. 2011. Bacterial transcription terminators: the RNA 3’-end chronicles. J Mol Biol, 412, 793–813.

Rappsilber, J., Ishihama, Y. & Mann, M. 2003. Stop and go extraction tips for matrix-assisted laser desorption/ionization, nanoelectrospray, and LC/MS sample pretreatment in proteomics. Anal Chem, 75, 663–70.

Sberro, H., Fremin, B. J., Zlitni, S., Edfors, F., Greenfield, N., Snyder, M. P., Pavlopoulos, G. A., Kyrpides, N. C. & Bhatt, A. S. 2019. Large-Scale Analyses of Human Microbiomes Reveal Thousands of Small, Novel Genes. Cell, 178, 1245–1259 e14.

Shell, S. S., Wang, J., Lapierre, P., Mir, M., Chase, M. R., Pyle, M. M., Gawande, R., Ahmad, R., Sarracino, D. A., Ioerger, T. R., Fortune, S. M., Derbyshire, K. M., Wade, J. T. & Gray, T. A. 2015. Leaderless Transcripts and Small Proteins Are Common Features of the Mycobacterial Translational Landscape. PLoS Genet, 11, e1005641.

Smith, C., Canestrari, J. G., Wang, J., Derbyshire, K. M., Gray, T. A. & Wade, J. T. 2019. Pervasive Translation in <em>Mycobacterium tuberculosis</em>. bioRxiv, 665208.

Stojiljkovic, I., Baumler, A. J. & Hantke, K. 1994. Fur regulon in gram-negative bacteria. Identification and characterization of new iron-regulated Escherichia coli genes by a fur titration assay. J Mol Biol, 236, 531–45.

Swaminathan, S., Ellis, H. M., Waters, L. S., Yu, D., Lee, E. C., Court, D. L. & Sharan, S. K. 2001. Rapid engineering of bacterial artificial chromosomes using oligonucleotides. Genesis, 29, 14–21.

Turnbough, C. L., Jr. 2019. Regulation of Bacterial Gene Expression by Transcription Attenuation. Microbiol Mol Biol Rev, 83.

Van Kessel, J. C. & Hatfull, G. F. 2008. Efficient point mutagenesis in mycobacteria using single-stranded DNA recombineering: characterization of antimycobacterial drug targets. Mol Microbiol, 67, 1094–107.

Weaver, J., Mohammad, F., Buskirk, A. R. & Storz, G. 2019. Identifying Small Proteins by Ribosome Profiling with Stalled Initiation Complexes. MBio, 10.

Xu, X., Vilcheze, C., Av-Gay, Y., Gomez-Velasco, A. & Jacobs, W. R., Jr. 2011. Precise null deletion mutations of the mycothiol synthesis genes reveal their role in isoniazid and ethionamide resistance in Mycobacterium smegmatis. Antimicrob Agents Chemother, 55, 3133–9.

Yanofsky, C. 1981. Attenuation in the control of expression of bacterial operons. Nature, 289, 751–8.

Zhao, G., Weatherspoon, N., Kong, W., Curtiss, R., 3RD & Shi, Y. 2008. A dual-signal regulatory circuit activates transcription of a set of divergent operons in Salmonella typhimurium. Proc Natl Acad Sci U S A, 105, 20924–9.

Zheng, X., Hu, G. Q., She, Z. S. & Zhu, H. 2011. Leaderless genes in bacteria: clue to the evolution of translation initiation mechanisms in prokaryotes. BMC genomics, 12, 361.

Zougman, A., Selby, P. J. & Banks, R. E. 2014. Suspension trapping (STrap) sample preparation method for bottom-up proteomics analysis. Proteomics, 14, 1006–0.

