## Supplementary Information for "Polycysteine-encoding leaderless short ORFs function as cysteine-responsive attenuators of operonic gene expression in mycobacteria"

### Supplementary Information file for Canestrari et al.

Running title: *Attenuating sORFs coordinate a multilocus regulon*

Table of Contents

| Page | Item | Description |
|---|---|---|
| 1. | Table of Contents | |
| 2. | Fig S1 | Expression profiles of *Ms5788A* and *Rv0815A* loci |
| 3. | Fig S2 | Predicted *Ms5788* leader mRNA folded structure and annotations |
| 4. | Fig S3 | Clustered mutations showing alternative G:U base pairing |
| 5. | Fig S4A, B | Expression profiles of *Ms0113A* and *Ms0932A* loci |
| 6. | Fig S4C, D | Expression profiles of *Ms4527A* and *Ms4333A* loci |
| 7. | Fig S4E, F | Expression profiles of *Ms4536A* and *Ms5280A* loci |
| 8. | Fig S4G | Expression profiles of the *Rv2334* locus |
| 9. | Table S1 | LL-sORFs identified in *Mycobacterium smegmatis* (1-76) |
| 10. | Table S1 | LL-sORFs identified in *Mycobacterium smegmatis* (77-145) |
| 11. | Table S1 | LL-sORFs identified in *Mycobacterium smegmatis* (146-230) |
| 12. | Table S1 | LL-sORFs identified in *Mycobacterium smegmatis* (231-304) |
| 13. | Table S2 | Amino acid distribution frequencies in annotated genes and LL-sORFs |
| 14. | Table S3 | Clustered cysteine total of each LL-sORF of each species |

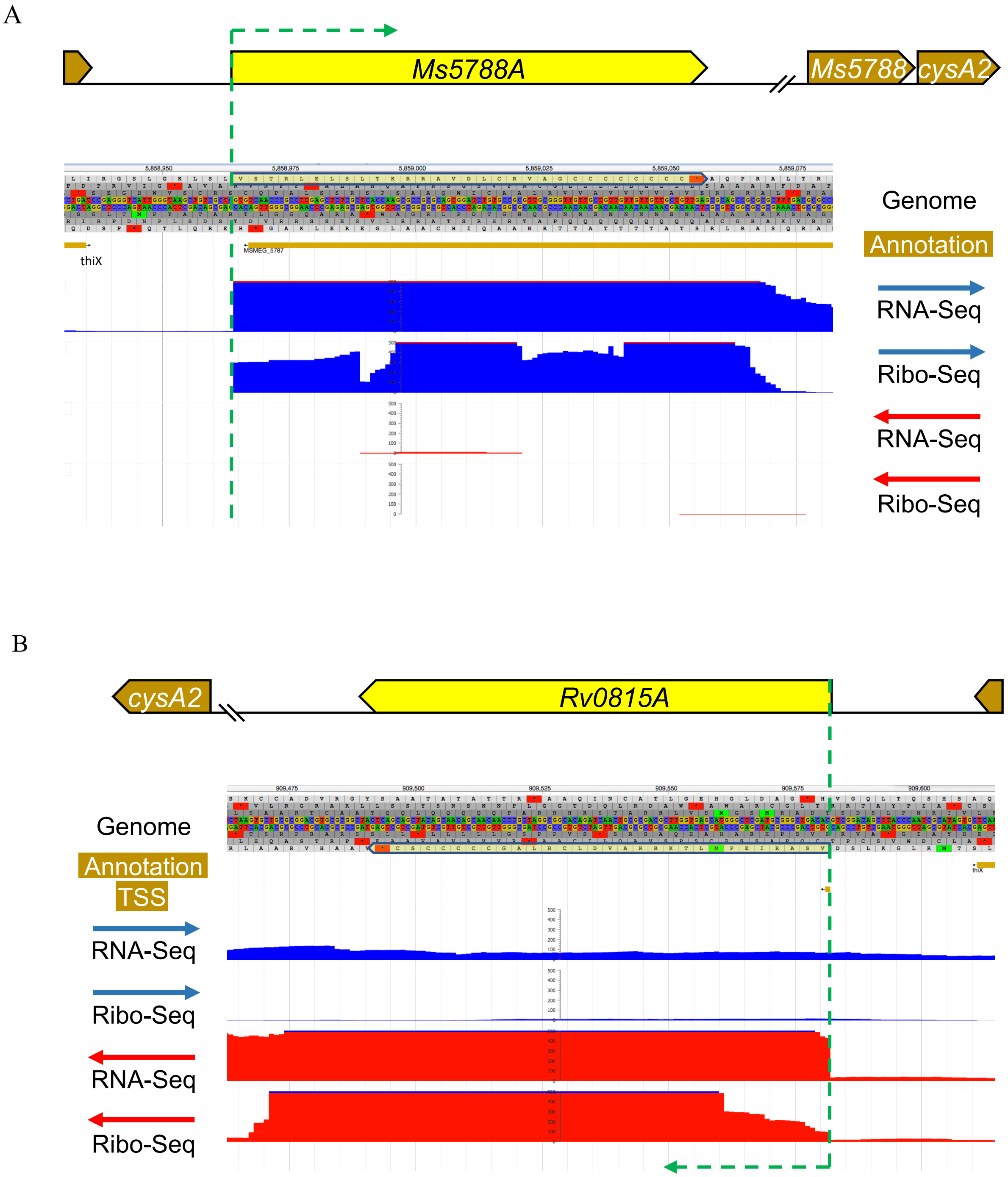

Supporting Information Fig S1. A. Schematic and expression profile of the *Ms5788A* locus. RNA-seq and Ribo-seq profiles show robust occupancy of the positive strand (maximum displayed read depth = 500), corresponding to the LL-sORF (shaded yellow). The end of the transcriptionally independent *thiX* gene is shown, and *cysA2* (Ms5789) represents operon genes downstream. Reads from a single replicate are shown for space efficiency. Note that *Ms5787* (gold line shown as an annotated gene reading from right to left) is likely a misannotation, as we detect no transcriptional or translational activity for this anti-sense gene. B. Schematic and expression profile for the *M. tuberculosis* orthologous locus, *Rv0815A*.

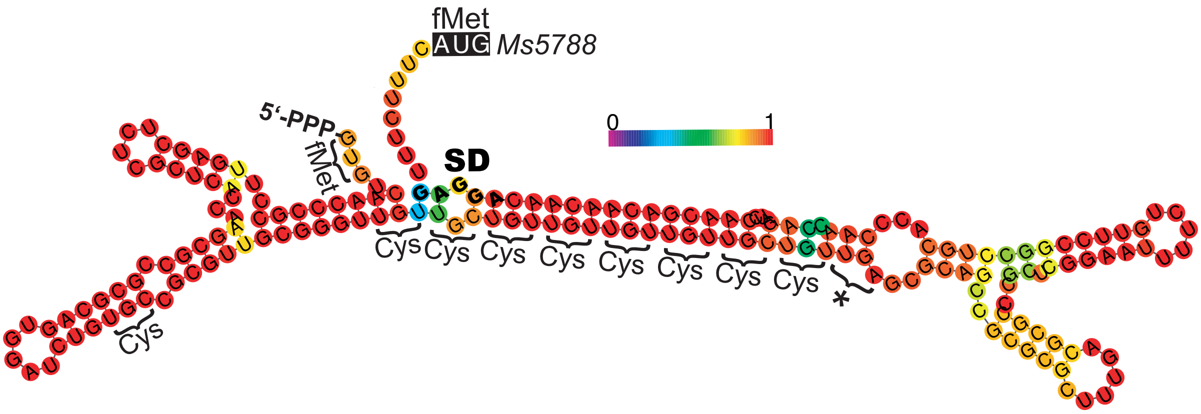

Supporting Information Fig S2. A stable mRNA structure model (RNAWebSuite/RNAfold) shows extensive base pairing forming a stem between the polycysteine coding region of the LL-sORF and the peri-SD region. The color of each nucleotide indicates the confidence in the predicted structure (as indicated in the heatmap gradient key).

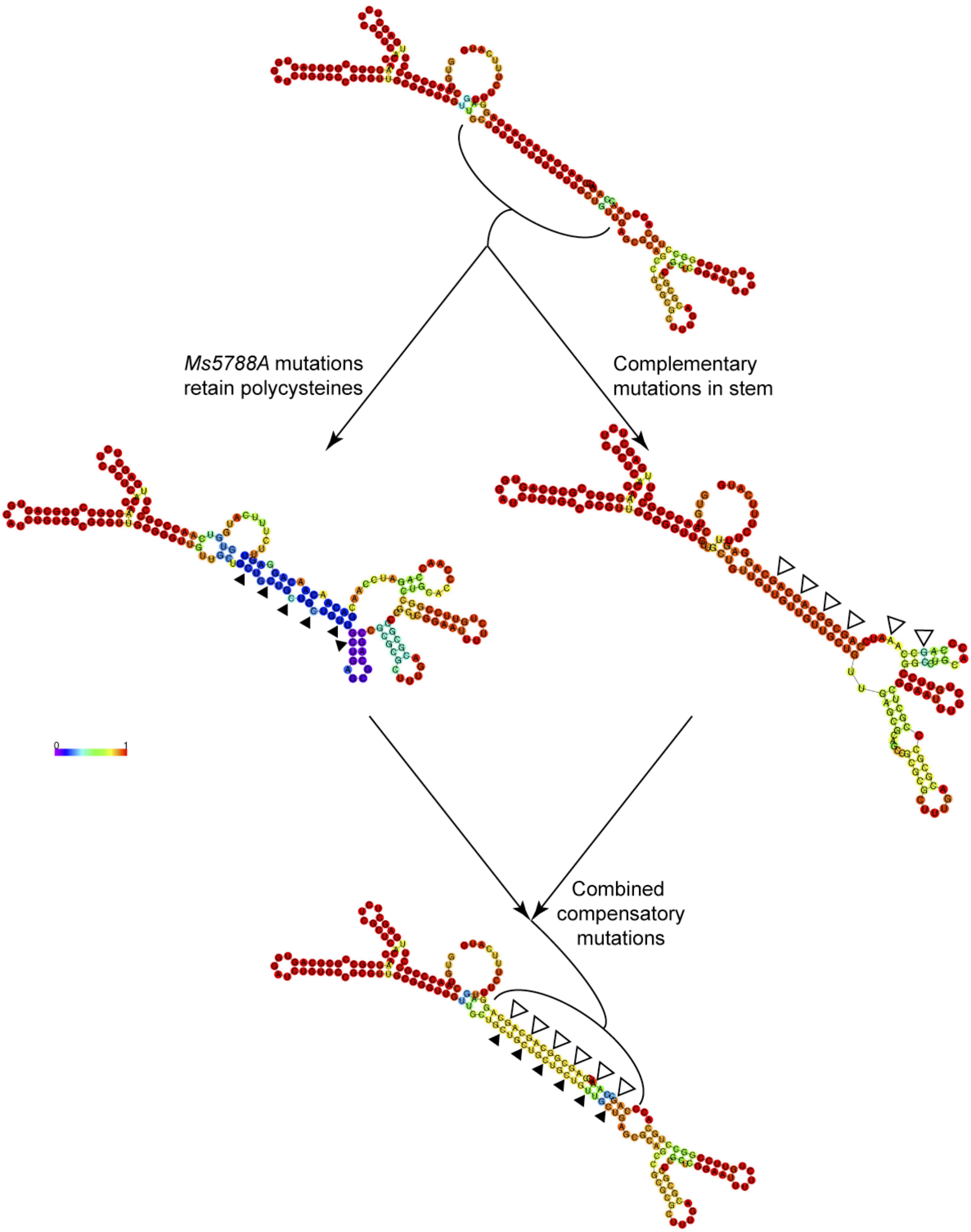

Supporting Information Fig S3. RNA-folding model of the polycysteine *Ms5788A* mutant. The *Ms5788A* ORF mutations (indicated by filled triangles) retain polycysteine coding but disrupt the stability and folding of the mRNA as indicated in the modeled structures. Mutations of the complementary nucleotides (indicated by open triangles) near the *Ms5788* SD sequence, are predicted to form G-U base pairs; the resulting overall structure and stability are comparable to that of the wild-type mRNA and delivers comparable expression of the reporter (Fig 4 vi).

4A

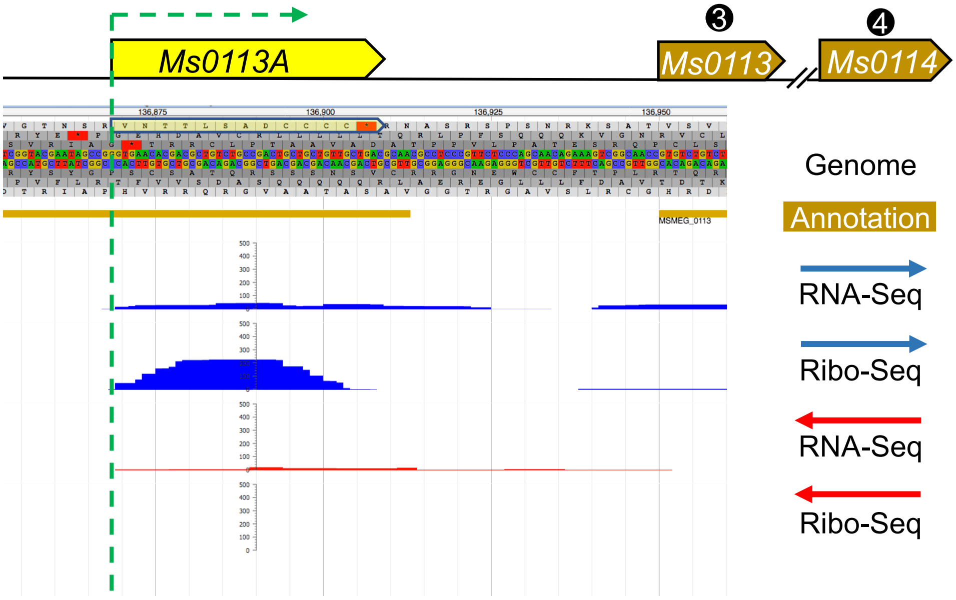

4B

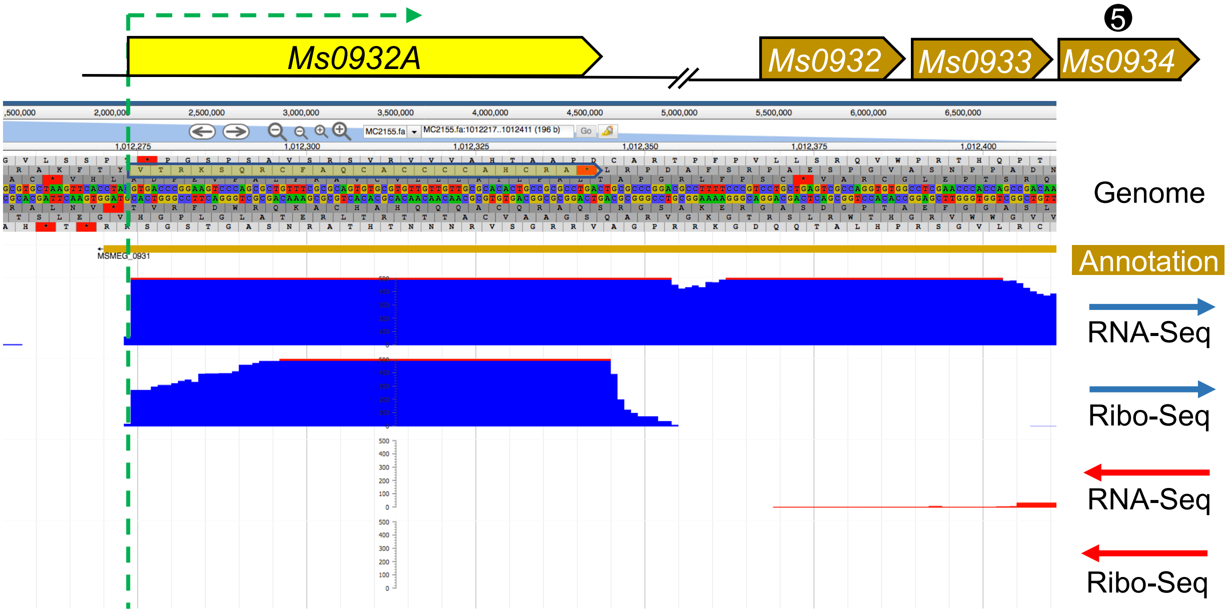

Supporting Information Fig S4. Additional loci that feature expressed polycysteine LL-sORFs upstream of annotated genes. JBrowse images of each LL-sORF locus show the gene context and sequence and transcriptomic profiles. Yellow arrow boxes indicate the LL-sORF in each locus, and the gene encoding the cysteine-responsive protein detected by mass spectrometry is indicated by the black circle with white number as in the mass spectrometry scatter plot (Fig 5a). Panels are shown for A) *Ms0113A/0013/Ms0114*, (B) *Ms0932A/0932/0933/0934*. Note that genes (B) *Ms0931* and (C) *Ms4528* are likely misannotated based on our profiling data. Pipeline prediction algorithms annotated none of these LL-sORFs. Screen shots captured from http://www.wadsworth.org/research/scientific-resources/interactive-genomics.

4C

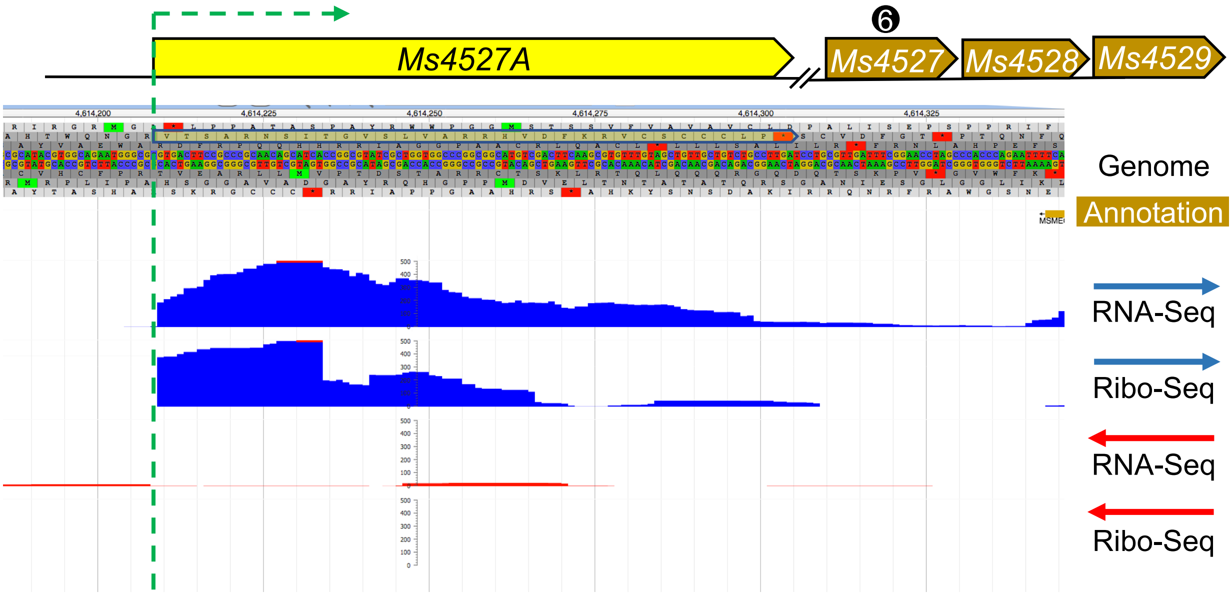

4D

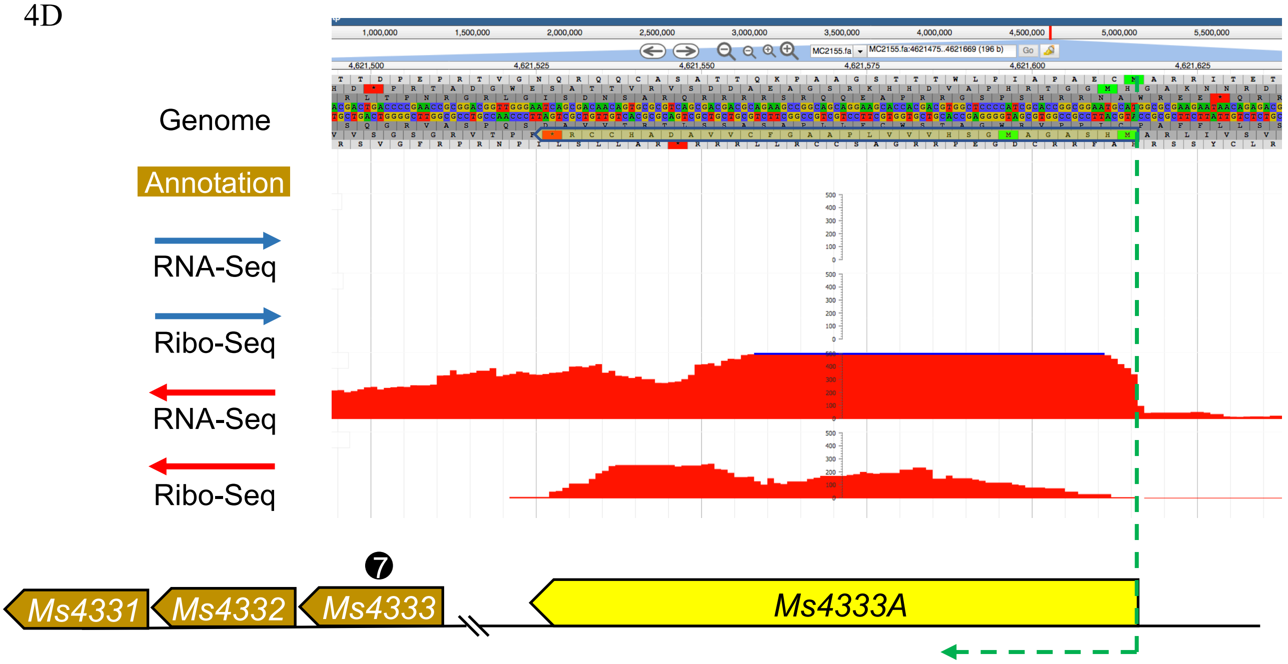

(C) *Ms4527A/4527/4528/4529*, (D) *Ms4533A/4533/4532/4531/4530*

4E

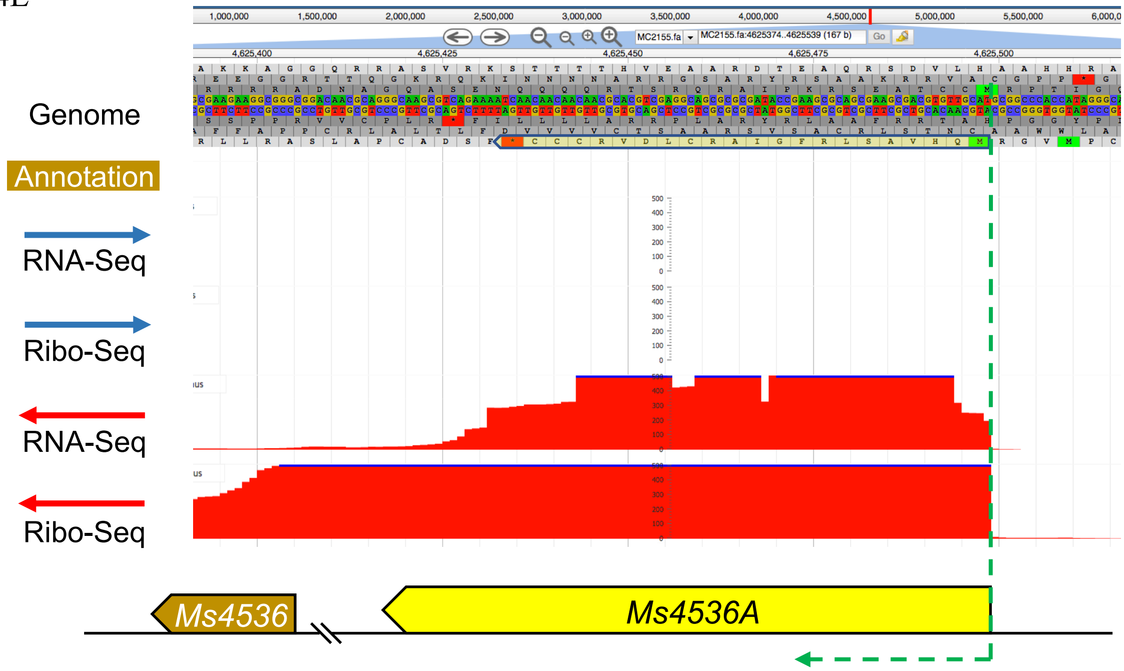

4F

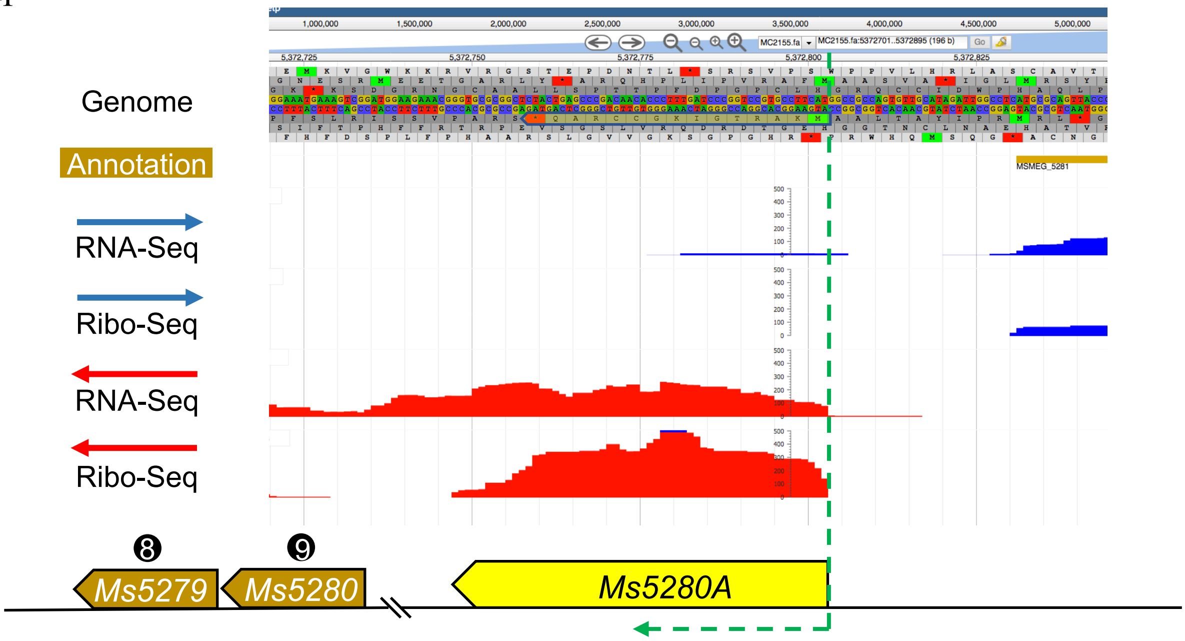

E) *Ms4536A/4536*, (F) *Ms5280/5279*

4G

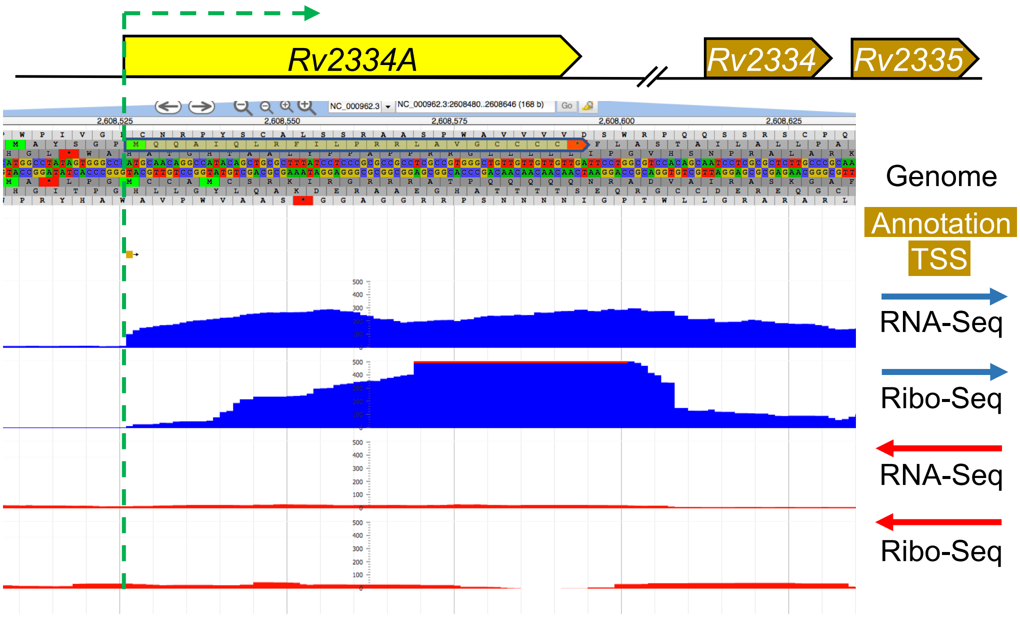

Rv2334A
Rv2334
Rv2335
Genome
Annotation
TSS
RNA-Seq
Ribo-Seq
RNA-Seq
Ribo-Seq

(G) *Rv2334A*

### Supporting Information Table S1. LL-sORF sequences identified in *Mycobacterium smegmatis*

| Coordinate | Strand | Amino Acid Sequence | Length | Downstream Gene | Mtb Ortholog |
|---|---|---|---|---|---|
| 3308801 | + | M- | 2 | MSMEG_3227 | Rv1617 |
| 4344259 | + | M- | 2 | MSMEG_4260 | Rv2193 |
| 1680101 | - | M- | 2 | MSMEG_1589 | |
| 2082138 | - | M- | 2 | MSMEG_2000 | |
| 6687146 | - | V- | 2 | MSMEG_6636 | |
| 3725020 | - | M- | 2 | Intragenic Sense | |
| 1776216 | + | M- | 2 | Intragenic Antisense | |
| 3315237 | + | V- | 2 | Intragenic Antisense | |
| 1340338 | - | V- | 2 | Intragenic Antisense | |
| 4227840 | - | V- | 2 | Intragenic Antisense | |
| 6722887 | - | M- | 2 | Intragenic Antisense | |
| 4320835 | - | VG- | 3 | MSMEG_4236 | Rv2166c |
| 6267182 | - | VA- | 3 | MSMEG_6199 | Rv3681c |
| 6476204 | - | VT- | 3 | MSMEG_6404 | Rv3809c |
| 4367370 | + | VC- | 3 | Intragenic Sense | |
| 2718889 | - | MT- | 3 | Intragenic Antisense | |
| 5046989 | - | MDR- | 4 | MSMEG_4951 | Rv1298 |
| 4764491 | + | MRA- | 4 | MSMEG_4676 | |
| 5455210 | - | MNR- | 4 | MSMEG_5375 | |
| 6687138 | - | MDS- | 4 | MSMEG_6636 | |
| 5637046 | + | VRM- | 4 | Intragenic Antisense | |
| 4812492 | + | MRWA- | 5 | MSMEG_4720 | Rv2507 |
| 1077518 | - | VRAA- | 5 | MSMEG_1007 | |
| 2370780 | - | VRAA- | 5 | MSMEG_2287 | |
| 5204785 | - | VLGI- | 5 | MSMEG_5102 | |
| 4069274 | + | VHHC- | 5 | Intragenic Antisense | |
| 943175 | + | MITM- | 5 | Intragenic Antisense | |
| 6442807 | - | VNAA- | 5 | Intragenic Antisense | |
| 1028988 | + | VVMKP- | 6 | MSMEG_0952 | Rv0509 |
| 1827558 | + | VGIAR- | 6 | MSMEG_1735 | Rv3303c |
| 3484156 | - | VRYGM- | 6 | MSMEG_3410 | |
| 3658192 | + | VTAER- | 6 | MSMEG_3595 | |
| 6476316 | + | MQSMY- | 6 | MSMEG_6405 | |
| 6602035 | - | VAVHG- | 6 | MSMEG_6544 | |
| 6731020 | - | MPLSI- | 6 | MSMEG_6677 | |
| 2279436 | + | VAIAN- | 6 | Intragenic Antisense | |
| 5551391 | - | MRWSQT- | 7 | MSMEG_5468 | Rv0996 |
| 5089826 | + | MAGRTT- | 7 | MSMEG_4995 | Rv1283c |
| 4562086 | - | VTLLCR- | 7 | MSMEG_4480 | |
| 3465554 | + | MPVRTR- | 7 | Intragenic Sense | |
| 691794 | + | MWARKG- | 7 | Intragenic Sense | |
| 5074019 | + | VRWTCR- | 7 | Intragenic Antisense | |
| 5673237 | + | VSDRRTC- | 8 | MSMEG_5584 | Rv0927c |
| 549810 | - | MPSLGES- | 8 | MSMEG_0472 | |
| 813287 | - | VEGVAWG- | 8 | MSMEG_0726 | |
| 2800986 | + | MTDLEAG- | 8 | MSMEG_2732 | |
| 3318373 | + | VAHFHGR- | 8 | MSMEG_3235 | |
| 3862543 | + | VSRDGRT- | 8 | MSMEG_3795 | |
| 4052066 | - | MTSEGQP- | 8 | MSMEG_3982 | |
| 4093727 | + | VTGKFAR- | 8 | MSMEG_4022 | |
| 3954589 | + | MRCRRFD- | 8 | Intragenic Sense | |
| 6704329 | + | VTKKKAE- | 8 | Intragenic Sense | |
| 4803609 | - | MTAQHAVT- | 9 | MSMEG_4712 | Rv2497c |
| 1726683 | + | VLCARRAG- | 9 | MSMEG_1634 | |
| 3787176 | + | MLRIVELG- | 9 | MSMEG_3718 | |
| 5565331 | - | VELLIRAT- | 9 | MSMEG_5483 | |
| 6032449 | - | VAEQSAGG- | 9 | MSMEG_5967 | |
| 6266602 | - | VVPDDQKQ- | 9 | MSMEG_6198 | |
| 6687148 | - | MCKDGQLA- | 9 | MSMEG_6636 | |
| 123288 | + | VVGVVVTN- | 9 | Intragenic Sense | |
| 4030320 | + | MQDAGVLT- | 9 | Intragenic Sense | |
| 5289435 | - | VDRTPACG- | 9 | Intragenic Sense | |
| 2267686 | + | VNWLRESG- | 9 | Intragenic Antisense | |
| 5285985 | + | VRTMKLPL- | 9 | Intragenic Antisense | |
| 6156790 | + | VSHSPSRS- | 9 | Intragenic Antisense | |
| 3767522 | - | MRQGGVLT- | 9 | Intragenic Antisense | |
| 4348093 | - | MQANVSSR- | 9 | Intragenic Antisense | |
| 1498103 | + | VPGPKGACG- | 10 | MSMEG_1398 | Rv0682 |
| 4787260 | - | VHQCRCAKA- | 10 | MSMEG_4699 | Rv2476c |
| 1507950 | + | VRASSSREL- | 10 | MSMEG_1407 | |
| 4457737 | - | MSAVMVASR- | 10 | MSMEG_4370 | |
| 4488056 | - | VCVDERYRL- | 10 | MSMEG_4396 | |
| 5151735 | + | VPISWIARA- | 10 | MSMEG_5051 | |
| 6547017 | - | MSRGFGVGA- | 10 | MSMEG_6484 | |
| 5633 | + | MTSVRVWRR- | 10 | Intragenic Sense | |
| 6784137 | + | MRTTPVGCR- | 10 | Intragenic Sense | |

| | | | | | |
|---|---|---|---|---|---|
| 1824107 | - | MTPSPGRER- | 10 | Intragenic Antisense | |
| 5349907 | + | MTHQLYVEVT- | 11 | MSMEG_5257 | Rv1085c |
| 3862355 | - | MDWVSSSDPT- | 11 | MSMEG_3793 | Rv1641 |
| 4356690 | - | VNARAIHAFR- | 11 | MSMEG_4272 | Rv2204c |
| 4734101 | - | VSSTAHRVCA- | 11 | MSMEG_4646 | Rv2455c |
| 5999994 | - | VIEQPPRSKK- | 11 | MSMEG_5940 | Rv3536c |
| 135742 | - | MPTDRLDRCR- | 11 | MSMEG_0110 | |
| 1832322 | - | MAVTPPKQGG- | 11 | MSMEG_1738 | |
| 5677010 | + | VVLIYLTVNG- | 11 | Intragenic Sense | |
| 775957 | + | MKSEKKAPLW- | 11 | Intragenic Antisense | |
| 5675072 | - | VAPRRPRVSR- | 11 | Intragenic Antisense | |
| 5044647 | - | MTDERSREESM- | 12 | MSMEG_4948 | Rv1301 |
| 6447255 | + | MNSVSEQQMTQ- | 12 | MSMEG_6388 | Rv3794 |
| 2185274 | - | MSSAGSGSGGA- | 12 | MSMEG_2106 | |
| 2333275 | + | MSGQRGCRCIV- | 12 | MSMEG_2251 | |
| 3601354 | + | VPTRSRRKGRS- | 12 | MSMEG_3540 | |
| 6304860 | - | VLGGSHDGVVR- | 12 | MSMEG_6239 | |
| 2088892 | + | VMMQSDVGGLR- | 12 | Intragenic Sense | |
| 3091687 | + | VAPYEGDLLHA- | 12 | Intragenic Sense | |
| 5363406 | + | VRSSPPLPTTV- | 12 | Intragenic Antisense | |
| 4350654 | - | MRAVRGRRRWR- | 12 | Intragenic Antisense | |
| 525025 | - | VRELNPTSGRF- | 12 | Intragenic Antisense | |
| 447603 | + | MFAMSAEQPGET- | 13 | MSMEG_0399 | Rv2377c |
| 2456666 | + | MAGKRRKFTHGS- | 13 | MSMEG_2374 | Rv3001c |
| 2840614 | + | MLPTGARHDRSA- | 13 | MSMEG_2775 | |
| 3511269 | - | VPSRESCGRPRH- | 13 | MSMEG_3441 | |
| 3906254 | - | MEPARSGIRRQP- | 13 | MSMEG_3836 | |
| 4048892 | - | VKGTGNRDATAP- | 13 | MSMEG_3979 | |
| 4212259 | + | MRIGEPLGVRVS- | 13 | MSMEG_4132 | |
| 4928308 | - | VRYVTQARIYSR- | 13 | MSMEG_4831 | |
| 4942300 | + | MLSRLDKHLSRP- | 13 | MSMEG_4847 | |
| 1535993 | + | MPVLGGAEGEVT- | 13 | Intragenic Sense | |
| 5791098 | + | MSSQPSSGVVSA- | 13 | Intragenic Sense | |
| 136869 | + | VNTTLSADCCCC- | 13 | Intragenic Antisense | |
| 5228464 | - | MPGVVDPECETRG- | 14 | MSMEG_5126 | Rv1173 |
| 6653443 | - | VKSSGKWGAGSCL- | 14 | MSMEG_6596 | Rv2172c |
| 6334571 | + | VQRVLLLGRRGGV- | 14 | MSMEG_6271 | Rv3710 |
| 1077583 | + | VLRRLVVATRSGV- | 14 | MSMEG_1008 | ** |
| 2192474 | - | VSAHNAAGRAMMR- | 14 | MSMEG_2116 | |
| 2370845 | + | VLRRLVVATRSGV- | 14 | MSMEG_2288 | ** |
| 2863607 | + | VTIGVCASYWRFI- | 14 | MSMEG_2799 | |
| 3759458 | + | MSSSRRVARSRIR- | 14 | MSMEG_3694 | |
| 4101893 | - | MGDTVRVDEAKVS- | 14 | MSMEG_4029 | |
| 4688794 | - | VELLSAATAGASR- | 14 | MSMEG_4598 | |
| 3756969 | + | VTPESVAQRLREV- | 14 | Intragenic Sense | |
| 5778189 | + | VRRGSRGAQSGLT- | 14 | Intragenic Sense | |
| 772378 | + | VPPASEVDAVHSG- | 14 | Intragenic Antisense | |
| 5372802 | - | MKARTGIKGCCRAQ- | 15 | MSMEG_5280 | Rv1065 |
| 5075326 | - | MLTILASSTNLMGK- | 15 | MSMEG_4979 | Rv1285 |
| 3211779 | + | MVGTGCPTPALGKL- | 15 | MSMEG_3138 | Rv1471 |
| 2817741 | - | MRGGARLPDRNARH- | 15 | MSMEG_2746 | Rv2714 |
| 76190 | + | MSGNRGDSCLTICG- | 15 | MSMEG_0056 | |
| 1053307 | + | MSGLWRHDALSHRR- | 15 | MSMEG_0981 | |
| 3357174 | + | VCRCFVRSYCSSGA- | 15 | MSMEG_3275 | |
| 406707 | + | VTDNPGLGPRVPVR- | 15 | Intragenic Sense | |
| 4433192 | + | VGMLIGLLLRRRRG- | 15 | Intragenic Sense | |
| 4126910 | - | VAVSSGSATRSDAV- | 15 | Intragenic Sense | |
| 4135536 | - | MCGLRRRACSTRIG- | 15 | Intragenic Sense | |
| 4868211 | - | MGVRSGRRGGRRVG- | 15 | Intragenic Sense | |
| 2949427 | - | VRSNVSIGMSSMIE- | 15 | Intragenic Antisense | |
| 3348749 | - | MRSRRRPRTGTGCR- | 15 | Intragenic Antisense | |
| 2400571 | - | VPRDVMRAADTRELD- | 16 | MSMEG_2317 | Rv3045 |
| 1107309 | - | VPRDVMRAADTRELD- | 16 | MSMEG_1037 | |
| 1257949 | - | MTAERKEGRSEEALE- | 16 | MSMEG_1191 | |
| 2480359 | - | MQRDGGDCGWSRRTC- | 16 | MSMEG_2397 | |
| 3519974 | + | MCSAYLKWNTFRIAT- | 16 | MSMEG_3452 | |
| 4943229 | + | MMPKTSLSLTPWVRR- | 16 | MSMEG_4849 | |
| 2596965 | + | VPIWYPKTARVITRR- | 16 | Intragenic Sense | |
| 4685015 | + | MGLALICHARVDSAR- | 16 | Intragenic Sense | |
| 2305056 | - | MDFDGGARCGKATGR- | 16 | Intragenic Sense | |
| 4873582 | - | MWSFSKSYPSMPRAR- | 16 | Intragenic Sense | |
| 117072 | + | VTPSRLHTSANLRPP- | 16 | Intragenic Antisense | |
| 2012567 | - | MSHWANMPQVRQSNTD- | 17 | MSMEG_1935 | Rv3208 |
| 6441234 | + | MSPRLGAAVDGVRRRR- | 17 | MSMEG_6385 | Rv3791 |
| 1395110 | + | MSDKWRARWIRTRIHR- | 17 | MSMEG_1304 | |
| 2098457 | + | VWVPCRRAMWSTARSC- | 17 | MSMEG_2020 | |
| 4688803 | - | MRVVELLSAATAGASR- | 17 | MSMEG_4598 | |

| | | | | | |
|---|---|---|---|---|---|
| 5684188 | - | MSAATVIAPSTNGAPR- | 17 | MSMEG_5595 | |
| 1036817 | + | MSVRYSKYARQAYARA- | 17 | Intragenic Sense | |
| 2227913 | - | MAAAPPQPAGGLVPGS- | 17 | Intragenic Sense | |
| 415754 | + | MVDSRPVAPGLASAER- | 17 | Intragenic Antisense | |
| 1358293 | - | VVVHKSARSMRRHQPG- | 17 | Intragenic Antisense | |
| 5414354 | + | VNRPESARHIRNMARRS- | 18 | MSMEG_5328 | |
| 1003063 | + | MRSRLLTDTDRPEQAHT- | 18 | Intragenic Sense | |
| 1758040 | - | VSARSATSTTCCWSPTR- | 18 | Intragenic Sense | |
| 4782457 | - | VSRIWRPCRLRHVRSAA- | 18 | Intragenic Sense | |
| 5806079 | + | MVAIRSAPIPCGTKPMR- | 18 | Intragenic Antisense | |
| 5988445 | + | MITSTRQIATVAAIASV- | 18 | Intragenic Antisense | |
| 5578437 | - | VSISMSAGVRVRSARRS- | 18 | Intragenic Antisense | |
| 1439467 | + | MSKVKEQHGWPRRRRSPG- | 19 | MSMEG_1346 | Rv0640 |
| 5857566 | - | MNPEDHPTPASVFSRRDS- | 19 | MSMEG_5784 | Rv0818 |
| 447513 | + | VHCQLSGGVLVSSSSTKA- | 19 | MSMEG_0399 | Rv2377c |
| 2035922 | - | VTRPRTRLRCHEYGCRYG- | 19 | MSMEG_1957 | Rv3195 |
| 1617771 | - | VCPLILFFVAPGSSRLPR- | 19 | MSMEG_1528 | Rv3452 |
| 979951 | + | MHPPRRHPRRGTQGDDAP- | 19 | MSMEG_0895 | |
| 3976145 | + | VCGLGTRHTSTAVNLSVQ- | 19 | MSMEG_3903 | |
| 4583431 | + | MPPMPNPCSTGVSQVRQR- | 19 | Intragenic Antisense | |
| 5040245 | + | VPDERIGGRWARWNHPRG- | 19 | Intragenic Antisense | |
| 3565763 | - | VLLSQKYAVRVSSPTEET- | 19 | Intragenic Antisense | |
| 553608 | - | VNPAACSGSTSISCVHPT- | 19 | Intragenic Antisense | |
| 4655982 | - | MSTKRSNRLPCHPAPISLV- | 20 | MSMEG_4564 | Rv2406c |
| 436171 | + | MAFREPQRRPFFCEEQRIS- | 20 | MSMEG_0387 | |
| 1742575 | - | MSITSMAAPVAAFIRPRTA- | 20 | MSMEG_1646 | |
| 6932359 | + | MPAADGAPSHGTMIERESR- | 20 | MSMEG_6884 | |
| 5625225 | + | VLIEQRRFFYGYRPAVPVV- | 20 | Intragenic Sense | * |
| 2551252 | - | VNASSGRAPRARDRSRTDR- | 20 | Intragenic Antisense | |
| 6597298 | - | VQYSAGLLRSNISATGSRS- | 20 | Intragenic Antisense | |
| 6654779 | - | VRRAVRASRQPRWRPRRPS- | 20 | Intragenic Antisense | |
| 6724109 | - | VCCRCPAEAAGVVPCRSVR- | 20 | Intragenic Antisense | |
| 2024947 | + | VRLLGSVQANTGRLSWMTRG- | 21 | MSMEG_1945 | Rv3200c |
| 1307974 | + | VRQLIFVLSTPCSADDGRAG- | 21 | MSMEG_1238 | |
| 2183134 | - | VTIVITEAFVPRESSIATHP- | 21 | MSMEG_2105 | |
| 5769986 | - | VWVTSSRPWPILLAGRFSTS- | 21 | MSMEG_5679 | |
| 1438060 | + | VWTLEGDLDEPVRYEEERSMR- | 22 | MSMEG_1344 | |
| 4145577 | - | VSAEESWVEVVPHDSRWAESY- | 22 | MSMEG_4071 | |
| 4625498 | - | MQHVASLRFGIARCLDVRCCC- | 22 | MSMEG_4536 | |
| 6311441 | - | MSSPEASKVGAFGHAVSPLNH- | 22 | MSMEG_6245 | |
| 6035128 | - | MSPVWATYSTFSRTSITSLCL- | 22 | Intragenic Sense | |
| 6146412 | + | VLSTLPQDLSSAFVHSALKTR- | 22 | Intragenic Antisense | |
| 5981025 | - | VAHAPVHRIRGRRRCGGHEAR- | 22 | Intragenic Antisense | |
| 6352896 | - | VNSPRAGRQQGSTGSGGCAGSP- | 23 | MSMEG_6286 | Rv3722c |
| 1322728 | + | VNFRLSIPAQYRSTLRSPGSRV- | 23 | MSMEG_1247 | |
| 1309947 | + | VDQNLPGFGAMEDAFMNWEDVW- | 23 | Intragenic Sense | |
| 1681086 | + | VTACLISEQRQARMSSNRRGGR- | 23 | Intragenic Sense | |
| 2051102 | - | VAKPAWMVKPCGSWPNCRGPAT- | 23 | Intragenic Sense | |
| 1012274 | + | VTRKSQRCFAQCACCCCAHCRA- | 23 | Intragenic Antisense | |
| 2082659 | + | VITELRSESDSEKPTDCHSVVE- | 23 | Intragenic Antisense | |
| 1324821 | - | VIPYEGRDGTTTVIRFGFTTSA- | 23 | Intragenic Antisense | |
| 4223159 | - | MVAASLSDPKVYSSSGSRLDRW- | 23 | Intragenic Antisense | |
| 2453995 | + | MQTPGVLVVIGWRVDAALALCRA- | 24 | MSMEG_2372 | Rv3003c |
| 3076240 | + | VIQIALKRRFDALRHVGSAAEFA- | 24 | MSMEG_3008 | |
| 4029403 | - | VSHGVHRSTRMLNCVPAQGMWVM- | 24 | MSMEG_3958 | |
| 5042661 | - | VLAEPEKPRGLPARPIGFRALRT- | 24 | MSMEG_4946 | |
| 6952557 | - | VTSAPISMRDVCSAHAPSMVQHS- | 24 | Intragenic Antisense | |
| 433179 | - | VLSHDQLKSYPGPAELDSRLTSAE- | 25 | MSMEG_0384 | Rv0334 |
| 5533231 | - | MVSVSRTGTNTELLIGEDAGTNRS- | 25 | MSMEG_5445 | Rv1003 |
| 1893939 | - | MTCAVYTIPHPSMRFRTLANRVLR- | 25 | MSMEG_1817 | Rv3277 |
| 1343050 | + | VNGWTLCPPDVNEIVEALKTNGVV- | 25 | MSMEG_1255 | |
| 5839545 | + | MPTSHSPFLSRPAELAALVAGYLT- | 25 | Intragenic Sense | |
| 5737248 | + | MQDCRGRREQGTRPPSEDGGRVHD- | 25 | Intragenic Antisense | |
| 5179640 | - | VPARIALMNRVIGMPHGRQTDRFEP- | 26 | MSMEG_5078 | Rv1213 |
| 1438100 | + | MKRNGVCGERRARRCRLRRRHGHGH- | 26 | MSMEG_1344 | |
| 890241 | + | VFRTLRWPTCCPHRRQTRSHRRTGE- | 26 | Intragenic Sense | |
| 3166485 | + | VVPSVSPMTAPRSRASRCGVRFPMK- | 26 | Intragenic Antisense | |
| 1334023 | + | VRTVMIMAAIRILLGSECADAQTQWR- | 27 | MSMEG_1253 | |
| 2090033 | - | VSESDHDHLIEPAMRQHDSISGTVHA- | 27 | MSMEG_2010 | |
| 6348005 | + | VTNSTSCTAVRSTTPSRRRTTRIHVC- | 27 | Intragenic Sense | |
| 4873615 | - | VSAWRDIRCPGMWSFSKSYPSMPRAR- | 27 | Intragenic Sense | |
| 2329886 | - | VTTIPHSVTQPATTRWLLNSREFTTR- | 27 | Intragenic Antisense | |
| 3002164 | + | MPGRCGRVDGPRKLDRRSSTRKEGVPA- | 28 | MSMEG_2940 | Rv2603c |
| 1942002 | - | VAPSPRSDPTWLKLVCTVGADLDEDCS- | 28 | MSMEG_1864 | |
| 6519962 | - | VRALRYRGVTGTAITTSQETAPCISIR- | 28 | MSMEG_6456 | |
| 1261045 | - | VSTFTPRRSAHIIGSAQSVAPNGRSAV- | 28 | Intragenic Antisense | |
| 4471237 | - | VKCRHSVAPAMVMPDPSTTWAVPRNIV- | 28 | Intragenic Antisense | |

| | | | | | | |
|---|---|---|---|---|---|---|
| 3318412 | + | VRAGSMVVNVSGLERGGTRPEAESRGNL- | 29 | MSMEG_3235 | | |
| 2274345 | + | VGRRTRCRLLWIRFRGVCRGDRSGRRRP- | 29 | Intragenic Sense | | |
| 1339860 | - | VKSAFGAVDESTTPQGRGRNMQPLADCH- | 29 | Intragenic Antisense | | |
| 5279036 | - | MLWEERTCCNESQPRRLPRRAASSWWPRS- | 30 | MSMEG_5178 | Rv1146 | |
| 4621615 | - | MHSAGAMGSHVVVLPAAGFCVVADAHCCR- | 30 | MSMEG_4533 | Rv2400c | |
| 6098692 | + | MLITPRTERSRVVVGSHRLARKDSRWPAW- | 30 | MSMEG_6032 | | |
| 6819732 | - | MVASGRDTGGRTGPALRARARIRTSTRGD- | 30 | MSMEG_6769 | | |
| 339787 | + | MDEKGLNREDRLLHRFEPRRLHRRREPES- | 30 | Intragenic Sense | | |
| 6947698 | + | MPKVISGSSTITATAIAEPRRTHFQLMRN- | 30 | Intragenic Antisense | | |
| 273030 | - | VQVRSRSDTPGISGHRACRRESGDFASSSS- | 31 | MSMEG_0241 | Rv0202c | |
| 1324918 | + | VRRGLLGYTRHPGWLDGWMAGWLDGCGVCT- | 31 | MSMEG_1251 | | |
| 5858964 | + | VSTRLELSLTKRRAVDLCRVAGCCCCCCCC- | 31 | MSMEG_5788 | | * |
| 4401804 | + | MSLAKGLSDGRFEQCCSLIPHPQPRRGDRA- | 31 | Intragenic Sense | | |
| 4532688 | + | VRRDQRYMTRGRIHAEVADLLDRTVAMNGE- | 31 | Intragenic Sense | | |
| 6869085 | - | MGWDCWRTPWSPKCDPTPWCSTTARCGAAT- | 31 | Intragenic Sense | | |
| 4374673 | + | VRMSGRRVPRRDAVHTDVCWSSPADTLKPIA- | 32 | MSMEG_4290 | Rv2220 | |
| 4614209 | + | VTSARNSITGVSLVARRHVDFKRVCSCCCLP- | 32 | MSMEG_4527 | Rv2391 | |
| 4207184 | + | MPTRGSTRATQTLVRSDSRRGPRQLQPRVRQ- | 32 | MSMEG_4126 | | |
| 5633376 | + | VSVFEVMHEVYPAFTHVETHNRRTMRSAWCT- | 32 | MSMEG_5547 | | |
| 5456135 | + | MPQLAQTTRSVWDVLRRRNGRWRNPEMVSQL- | 32 | Intragenic Antisense | | |
| 4988791 | + | MWTSEGGWWRGFAGEPSLRYVRSRWPGAAATR- | 33 | MSMEG_4893 | Rv0419 | |
| 5860943 | + | VVLFFEILLVAAVLVITWFAVYALYRLVTDES- | 33 | Intragenic Sense | | |
| 1814971 | - | VEWVCWIGSGRFGCIGDRFRLRVGRRWRGVRIG- | 34 | MSMEG_1717 | | |
| 2751933 | + | VEWVCWIGSGRFGCIGDRFRLRVGRRWRGVRIG- | 34 | MSMEG_2675 | | |
| 153326 | - | VKMMSTVSGIDTASVGRKATRMRNQLWRTNSFH- | 34 | Intragenic Antisense | | |
| 3520180 | - | MPTPASLAMAAGAVSSPCSANALAATSRMRARF- | 34 | Intragenic Antisense | | |
| 193272 | - | VIVVTCCPTDRLVTRHESVCNIQNPQVRPLPADP- | 35 | MSMEG_0167 | | |
| 3169751 | - | MPQARRKSELRKSPIRNRSRGCHGTVFHRLLRAR- | 35 | MSMEG_3095 | | |
| 6413180 | + | MRQWRHGFAVELWGSGRCSRARFHVSFPRCPRRC- | 35 | MSMEG_6354 | | |
| 1808404 | + | VTQTTAFRSGGTWRQEPVFSGVIVRCRQTEQRGLR- | 36 | MSMEG_1713 | | |
| 4265402 | - | VTASDTSTHRTATCVRAMRRGCCMRRRHDGLTVPC- | 36 | MSMEG_4185 | | |
| 2432524 | + | MAGRSPASSRRTNQSHDEHRGPDQTGPRHMVRAQAL- | 37 | MSMEG_2351 | Rv3029c | |
| 141850 | - | MDPNPDYDLSDEGEFFFNWIPWGLRGVYPPPAYPPV- | 37 | MSMEG_0119 | | |
| 1523567 | - | MRSSSSMMSLHCCVDNSRGSRLPDEDRQWTHARLHA- | 37 | Intragenic Antisense | | |
| 1817937 | + | MHTASTTLTTRHRRICAATRSSRRSHQDGASLRKATR- | 38 | MSMEG_1723 | | |
| 6194162 | - | VLSLKQRVSATRVAIVVKDVIFPCARHLTESLRRPLT- | 38 | MSMEG_6131 | | |
| 6488567 | - | VRYLGSWPTESAAGAPPPRLDEATTWLEGLRAGSGGA- | 38 | Intragenic Sense | | |
| 5497774 | - | VVDLSDVMGASGSTIGSEDAGEEARPGFRKSASTTTM- | 38 | Intragenic Antisense | | |
| 6420693 | + | VRCGFGVLMGLLLPDHPPRSSPGRRAVHTGGDGSGRLP- | 39 | MSMEG_6364 | | |
| 305260 | - | VQMIGAAGRLYIGGSTDEVTVARDHITTVLSAIEGQEH- | 39 | Intragenic Sense | | |
| 6309094 | - | MRTRHASINTAPIAESERKFHWPLGRRPDDHPSLSDLGL- | 40 | Intragenic Sense | | |
| 1253931 | - | MSDVAERAHCRRTLPTSVLSFASPHKQRIGTRDREGCAVA- | 41 | MSMEG_1188 | | |
| 4240389 | - | VLPKSTVPNYRRPTLALYARRSIWRPQIRQGQNGPSNLRQ- | 41 | MSMEG_4160 | | |
| 4734022 | - | VTKTTVLSRTRLQHVSELGIPVCCARVSWGRAVRSSLRAMQ- | 42 | MSMEG_4646 | Rv2455c | |
| 4153805 | - | VGDMSAQTVGDDPVWPPRFRERREASSGRTWLLRTRDGATA- | 42 | MSMEG_4079 | | |
| 5193214 | + | VPHACEEGARRWVRLTDSIRRGFLATASISNGRESAPWQQR- | 42 | MSMEG_5093 | | |
| 5781120 | + | VLILGLAQTIVSYRGESLIPGFGGNRPVLAEREGTMAWGAR- | 42 | Intragenic Sense | | |
| 524687 | + | VPFGAEAATAEPMTSSTAYAIIMRRWGRCCANCTASTVPTA- | 42 | Intragenic Antisense | | |
| 1797855 | + | VAPLRRIDAQVDLAGRAARGAGHGDDPVRMWTVRRRWRRWRR- | 43 | MSMEG_1704 | | |
| 496059 | + | VSRNAKRNGSEMNAIGMNHGRSPTHWPKWLDPGAHAVLTHGE- | 43 | Intragenic Antisense | | |
| 107453 | - | VRSARRLGMRRCLSAQTPICLLGGSGPRRSDQFLLGECEACSA- | 44 | MSMEG_0083 | Rv3883c | |
| 3482565 | - | MRPPPDLLPDGTAGWAPVTAGESGAAVLRDHSGQRASTCYSIL- | 44 | MSMEG_3407 | | |
| 5279899 | + | MWIDDTSADVIKVDFEALYHGDVLVEGDTSEQFDDFQQLPTAV- | 44 | Intragenic Sense | | * |
| 6467259 | - | VPHELDLPGEVHRGRWQERRVQLPAERYAQLGVLGPAAAGDEA- | 44 | Intragenic Sense | | |
| 1928531 | - | MGQGGVSGLVTGAVFKIAERQFLPLAGSIPVRLRHRMPGPLLHD- | 45 | MSMEG_1851 | | |
| 892020 | + | VDRDALQIHGEGVAFTVGLYSQDQRGRNGIPVRDDRTRHQISPQ- | 45 | Intragenic Sense | | |
| 4693186 | + | VLAQLFDGEFGALCDRLFGRALRGDHGRGAEHKWRRCPRLGCRA- | 45 | Intragenic Antisense | | |
| 1371260 | - | VHLLQRCDDQQLEPESCEEIERVPCRHVRSAPERLVDDGEAEVA- | 45 | Intragenic Antisense | | |
| 5284781 | - | MSAVNRSTREGAFEHGRTTARFAGCRVGRYRPRPARGDDSGRSGR- | 46 | MSMEG_5184 | Rv1143 | |
| 4699300 | - | MTPDAGLSERWRTSDGFGGRHRCNHNGAVPNRSARQCVPGCTRRR- | 46 | MSMEG_4614 | | |
| 3816915 | - | VTASPLPGPSGNVEYFLRLRAETDSPLEGDALEAAVRRAVEEGPQ- | 46 | Intragenic Sense | | |
| 6239215 | + | VSRHRAQKRRQERRLACAGSAGDQESEPGVDDPAQQIDRGRRDRA- | 46 | Intragenic Antisense | | |
| 1458621 | + | VTVEAERPIPITRRHAWPRLTRPRRLPTLPSSSRRRRPPSSPSTAG- | 47 | MSMEG_1364 | Rv0651 | |
| 3811745 | - | VGWQATASLSDSLPFVTEDAFARVTQAPANRHETPLRHRWSCFVAG- | 47 | MSMEG_3746 | Rv1699 | |
| 4692797 | - | MARTPLHMFGMEYRRDEYLGLVASGEHDVNAQDLDGKTPLHYATEQ- | 47 | MSMEG_4605 | | |
| 4137551 | - | MSCCGPVVMLCVCRAGVRRLCPGWTLRVSSIRSGRAVMVAFMWGSG- | 47 | Intragenic Sense | | |
| 5306665 | + | VVSPEGVAEFLEKVPAAEFVELSDAGHTAAGDDNDAFTQVVVEFVNR- | 48 | Intragenic Sense | | |
| 5650281 | + | VAAPAGRVRAPPGRVVLGEGALGHRRQRAGRRTRDPRTAGGSGPHHQ- | 48 | Intragenic Sense | | |
| 3453984 | - | MSTAFWIVIAILAIGTIAGGLLRMRAWLQRPVPPEVIEAAHRNDDEDG- | 49 | MSMEG_3383 | | |
| 2407176 | + | MRKNWLSRLSVKYRRAYQPHPRTSARDADARRVHSELDAIRARFPDHA- | 49 | Intragenic Sense | | |
| 5375958 | + | VRCCDCWTRRCRQRGSCGPRRSSTRCATSSETSPSRSCPSLTPRSRPI- | 49 | Intragenic Sense | | |
| 5773222 | + | MSTYWVSSLTSMLRRQRLIILVRAMLQTSPSRLRFTSTQVLYNGTAGIE- | 50 | Intragenic Antisense | | |
| 6276556 | - | MPRGDGIYDEEHRDHDTGSAAEGGAGKPKRVPEDTPDVEMPDETTQEPPD- | 51 | MSMEG_6211 | | |
| 6549829 | - | VSYQAGETRHFHFGPGEFLLHDIENIGDTELVFTTVEHLDSDNAPLPVTR- | 51 | Intragenic Sense | | |

| ANNOTATED | | | | | | | | | | | | | | | | | | | | Total All | Total Poly | %Poly |
|---|---|---|---|---|---|---|---|---|---|---|---|---|---|---|---|---|---|---|---|---|---|---|
| Amino Acid | x1 | x2 | x3 | x4 | x5 | x6 | x7 | x8 | x9 | x10 | x11 | x12 | x13 | x14 | x15 | x16 | x17 | x18 | x19 | | | |
| A, alanine | 197695 | 29314 | 4605 | 729 | 186 | 33 | 1 | 1 | 1 | | | | | | | | | | | 274206 | 76511 | 27.9 |
| C, cysteine | 16898 | 197 | 1 | 1 | | | | | | | | | | | | | | | | 17299 | 401 | 2.318 |
| D, aspartate | 115764 | 7902 | 627 | 26 | 4 | | 6 | | | | | | | | | | | | | 133615 | 17851 | 13.36 |
| E, glutamate | 103393 | 5090 | 167 | 6 | | | | | | | | | | | | | | | | 114098 | 10705 | 9.382 |
| F, phenylalanine | 61161 | 2052 | 73 | 1 | | 1 | | | | | | | | | | | | | | 65494 | 4333 | 6.616 |
| G, glycine | 155845 | 13140 | 1222 | 123 | 25 | 5 | 2 | | | 1 | | | 1 | | | | | | | 186475 | 30630 | 16.43 |
| H, histidine | 45231 | 1405 | 54 | 2 | 1 | | | | | | | | | | | | | | | 48216 | 2985 | 6.191 |
| I, isoleucine | 85428 | 3059 | 82 | 1 | | | | | | | | | | | | | | | | 91796 | 6368 | 6.937 |
| K, lysine | 43100 | 1377 | 37 | 3 | | | | | | | | | | | | | | | | 45977 | 2877 | 6.257 |
| L, leucine | 168666 | 16936 | 1363 | 158 | 9 | 1 | 1 | | | | | | | | | | | | | 207317 | 38651 | 18.64 |
| M, methioine | 41595 | 881 | 16 | 3 | | | | | | | | | | | | | | | | 43417 | 1822 | 4.197 |
| N, asparagine | 44477 | 1187 | 33 | | | | | | | | | | | | | | | | | 46950 | 2473 | 5.267 |
| P, proline | 107885 | 5629 | 544 | 103 | 33 | 19 | 11 | 2 | 3 | 1 | | | | 1 | | | | | | 121610 | 13725 | 11.29 |
| Q, glutamine | 56801 | 2368 | 115 | 5 | 3 | | 1 | | | | | | | | | | | | | 61924 | 5123 | 8.273 |
| R, arginine | 127562 | 11444 | 1220 | 138 | 15 | 6 | 3 | | 1 | | | | | | | | | | | 154803 | 27241 | 17.6 |
| S, serine | 99161 | 5508 | 365 | 38 | 8 | 1 | 3 | | 1 | | | | | | | | | | | 111500 | 12339 | 11.07 |
| T, threonine | 114519 | 7025 | 569 | 42 | 10 | 4 | 4 | 1 | 2 | | 1 | | | | | | | | 1 | 130602 | 16083 | 12.31 |
| V, valine | 151768 | 14766 | 1590 | 156 | 10 | 1 | | | | | | | | | | | | | | 186750 | 34982 | 18.73 |
| W, tryptophan | 29995 | 632 | 10 | | | | | | | | | | | | | | | | | 31289 | 1294 | 4.136 |
| Y, tyrosine | 42884 | 1038 | 19 | 1 | | | | | | | | | | | | | | | | 45021 | 2137 | 4.747 |
| TOTALS | 1809828 | 130950 | 12712 | 1536 | 304 | 71 | 32 | 4 | 8 | 2 | 1 | 0 | 1 | 1 | 0 | 0 | 0 | 0 | 1 | 2118359 | 308531 | 14.56 |
| LL-sORFs | | | | | | | | | | | | | | | | | | | | | | |
| Amino Acid | x1 | x2 | x3 | x4 | x5 | x6 | x7 | x8 | x9 | x10 | x11 | x12 | x13 | x14 | x15 | x16 | x17 | x18 | x19 | Total All | Total Poly | %Poly |
| A, alanine | 437 | 40 | 3 | | | | | | | | | | | | | | | | | 526 | 89 | 16.92 |
| C, cysteine | 142 | 14 | 2 | 2 | | | | 1 | | | | | | | | | | | | 192 | 50 | 26.04 |
| D, aspartate | 204 | 15 | | | | | | | | | | | | | | | | | | 234 | 30 | 12.82 |
| E, glutamate | 212 | 11 | 1 | | | | | | | | | | | | | | | | | 237 | 25 | 10.55 |
| F, phenylalanine | 103 | 4 | 1 | | | | | | | | | | | | | | | | | 114 | 11 | 9.649 |
| G, glycine | 407 | 31 | | | | | | | | | | | | | | | | | | 469 | 62 | 13.22 |
| H, histidine | 159 | 2 | | | | | | | | | | | | | | | | | | 163 | 4 | 2.454 |
| I, isoleucine | 158 | 3 | | | | | | | | | | | | | | | | | | 164 | 6 | 3.659 |
| K, lysine | 88 | 2 | 1 | | | | | | | | | | | | | | | | | 95 | 7 | 7.368 |
| L, leucine | 285 | 27 | 3 | | | | | | | | | | | | | | | | | 348 | 63 | 18.1 |
| M, methioine | 90 | 4 | | | | | | | | | | | | | | | | | | 98 | 8 | 8.163 |
| N, asparagine | 101 | | | | | | | | | | | | | | | | | | | 101 | 0 | 0 |
| P, proline | 331 | 17 | 3 | | | | | | | | | | | | | | | | | 374 | 43 | 11.5 |
| Q, glutamine | 149 | 6 | | | | | | | | | | | | | | | | | | 161 | 12 | 7.453 |
| R, arginine | 571 | 95 | 12 | 4 | | | | | | | | | | | | | | | | 813 | 242 | 29.77 |
| S, serine | 419 | 35 | 5 | 3 | | | | | | | | | | | | | | | | 516 | 97 | 18.8 |
| T, threonine | 318 | 25 | 2 | | | | | | | | | | | | | | | | | 374 | 56 | 14.97 |
| V, valine | 283 | 27 | 4 | | | | | | | | | | | | | | | | | 349 | 66 | 18.91 |
| W, tryptophan | 114 | 2 | | | | | | | | | | | | | | | | | | 118 | 4 | 3.39 |
| Y, tyrosine | 63 | | | | | | | | | | | | | | | | | | | 63 | 0 | 0 |
| TOTALS | 4634 | 360 | 37 | 9 | 0 | 0 | 0 | 1 | 0 | 0 | 0 | 0 | 0 | 0 | 0 | 0 | 0 | 0 | 0 | 5509 | 875 | 15.88 |

Supporting Information Table S2. Amino acid distribution frequencies in annotated genes and LL-sORFs encoded in *Mycobacterium smegmatis*. Each amino acid is listed alphabetically by its single letter designation. The number of instances for single (x1) or consecutive (x2 or greater) appearances are shown in the body of the table. Tallies factor the size of the cluster, so that 6 instances of 7 consecutive aspartates totals 42 aspartates, as reflected in the total columns at right.

| LL-sORF | 0113A | 0932A | 4527A | 4533A | 4536A | 5280A | 5788A | Rv2334A |
|---|---|---|---|---|---|---|---|---|
| Mycobacterium_abscessus | 4 | | 3 | 2 | | 2 | 6 | |
| Mycobacterium_africanum | | 3 | 3 | 2 | | 2 | 6 | 4 |
| Mycobacterium_avium_104 | | | 3 | 2 | 6 | | 7 | |
| Mycobacterium_bovis | | 3 | 3 | 2 | | 2 | 6 | 4 |
| Mycobacterium_canettii | | 3 | 3 | 2 | | 2 | | 4 |
| Mycobacterium_caprae | | 3 | 3 | 2 | | 2 | 6 | 4 |
| Mycobacterium_chelonae | 3 | | 3 | 2 | | 2 | 6 | |
| Mycobacterium_chimaera | | | 3 | 2 | | | 7 | 3 |
| Mycobacterium_chubuense | | | 4 | 4 | | | 6 | |
| Mycobacterium_colombiense | | | 3 | 2 | 7 | | 7 | |
| Mycobacterium_dioxanotrophicus | | 3 | 4 | 2 | 4 | | 7 | |
| Mycobacterium_fortuitum | | 4 | 4 | 2 | | 2 | 7 | |
| Mycobacterium_gilvum | | | 4 | 3 | | | 7 | |
| Mycobacterium_goodii | 4 | 4 | 4 | 2 | | 2 | 7 | |
| Mycobacterium_haemophilum | | | 4 | | 5 | | 6 | |
| Mycobacterium_hassiacum | | | 4 | 3 | | | 6 | |
| Mycobacterium_immunogenum | 3 | | 3 | | | 2 | 7 | |
| Mycobacterium_intracellulare | | | 3 | 2 | | | 7 | 3 |
| Mycobacterium_kansasii | | 3 | | 2 | 5 | | | |
| Mycobacterium_leprae | | | | | 3 | | 3 | |
| Mycobacterium_lepraemurium | | | 4 | | 6 | | 5 | |
| Mycobacterium_liflandii | | 3 | 4 | | | | 6 | 5 |
| Mycobacterium_litorale | | | 4 | 2 | | | 6 | |
| Mycobacterium_marinum_M | | 3 | 4 | | | | 6 | 5 |
| Mycobacterium_marseillense | | | 3 | 2 | | | 7 | 3 |
| Mycobacterium_microti | | 3 | 3 | 2 | | 2 | 6 | 4 |
| Mycobacterium_neoaurum | 4 | | 4 | 3 | | 2 | 5 | |
| Mycobacterium_paragordonae | | 3 | 4 | 2 | 6 | | | |
| Mycobacterium_phlei | 3 | 4 | 4 | | | | 6 | |
| Mycobacterium_pseudoshottsii | | 3 | 4 | | | | 6 | 5 |
| Mycobacterium_rutilum | | 3 | 4 | 4 | | | 6 | |
| Mycobacterium_shigaense | | | 3 | 2 | 4 | 2 | 5 | |
| Mycobacterium_simiae | | | 3 | 2 | 4 | | 8 | |
| Mycobacterium_sinense | | | 3 | 2 | | 3 | 5 | |
| Mycobacterium_smegmatis | 4 | 4 | 4 | 2 | 3 | 2 | 8 | |
| Mycobacterium_stephanolepidis | 3 | | 3 | 2 | | 2 | 6 | |
| Mycobacterium_thermoresistibile | | 3 | 4 | | | | | |
| Mycobacterium_tuberculosis | | 3 | 3 | 2 | | 2 | 6 | 4 |
| Mycobacterium_ulcerans | | 3 | 4 | | | | | 5 |
| Mycobacterium_vaccae | | | 4 | 3 | | | 6 | |
| Mycobacterium_vanbaalenii | | 4 | 4 | 3 | | | 6 | |

Supporting Information Table S3. Clustered cysteine total of each LL-sORF in each species
